# Probing neuropeptide volume transmission in vivo by a novel all-optical approach

**DOI:** 10.1101/2021.09.10.459853

**Authors:** Hejian Xiong, Emre Lacin, Hui Ouyang, Aditi Naik, Xueqi Xu, Chen Xie, Jonghae Youn, Krutin Kumar, Tyler Kern, Erin Aisenberg, Daniel Kircher, Xiuying Li, Joseph A. Zasadzinski, Celine Mateo, David Kleinfeld, Sabina Hrabetova, Paul A. Slesinger, Zhenpeng Qin

## Abstract

Neuropeptides are essential signaling molecules in the nervous system involved in modulating neural circuits and behavior. Although hypothesized to signal *via* volume transmission through G-protein coupled receptors (GPCR), remarkably little is known about their extrasynaptic diffusion. Here, we developed an all-optical approach to probe neuropeptide volume transmission in mouse neocortex. To control neuropeptide release, we engineered photosensitive nanovesicles with somatostatin-14 (SST) that is released with near-infrared light stimulation. To detect SST, we created a new cell-based neurotransmitter fluorescent engineered reporter (CNiFER) using the SST2 GPCR. Under two-photon imaging, we determined the time to activate SST2R at defined distances as well as the maximal distance and loss rate for SST volume transmission in neocortex. Importantly, we determined that SST transmission is significantly faster in neocortex with a chemically degraded extracellular matrix, a diseased condition indicated in neuroinflammation and Parkinson’s disease. These new neurotechnologies can reveal important biological signaling processes previously not possible, and provide new opportunities to investigate volume transmission in the brain.

## Introduction

Neuropeptides comprise a diverse class of signaling molecules that regulate brain states, modify neural activity as well as control vascular tone^1, 2^. Neuropeptides are synthesized through protein translation, processed proteolytically, and then released from dense-core vesicles in peptide-expressing neurons^3^. In contrast to classical, fast synaptic transmission, neuropeptides diffuse from the release sites and exert their actions through G-protein coupled receptors (GPCRs) at relatively long distances. Furthermore, neuropeptides activate GPCRs at low nanomolar concentrations, up to 1000-fold lower compared with low affinity neurotransmitters for ionotropic receptors (micromolar for GABA or glutamate)^4^. This diffusion-driven distribution is referred to as volume transmission, a nonsynaptic dispersion in the extracellular space (ECS)^4^. Determining where a particular neuropeptide acts relative to its release site is critical to elucidating its functional role in controlling neural circuits. Several factors affect their rate and extent of diffusion, including degradation by extracellular peptidases and the tortuosity of the ECS in the brain^5–7^. These factors make it challenging to characterize neuropeptide and quantitatively measure the neuropeptide diffusion in the local brain environment. However, the current techniques available are inadequate to study the properties of neuropeptide released *in vivo*.

To overcome this deficiency in the field, we first need a tool to control neuropeptide release with high spatial and temporal resolution. Optical stimulation methods to trigger endogenous neuropeptide release from neurons are limited^8, 9^, and co-transmission is common^1^. Methods for direct peptide application, such as perfusion and pressure injection, offer poor control over the injected volume^10^. One promising approach is the use of caged compounds that can be photolyzed by ultraviolet, blue or two-photon light^11, 12^. However, caged compounds have some limitations such as limited tissue penetration of uncaging with ultraviolet light (≤ 50 μm depth) and the peptidase degradation once injected into the slice or *in vivo*^13, 14^ Recently, it has been demonstrated that biomolecules can be photoreleased rapidly within sub-second by using ultrashort picosecond^15^ or femtosecond^16^ near-infrared laser pulses to activate gold-coated plasmonic nanovesicles. The development of peptide-filled nanovesicles would provide an efficient and ultrafast release mechanism to study diffusion *in vivo*.

In addition to the fast temporal and spatial release, we need a highly sensitive detection method to measure neuropeptides *in vivo*. Recently, two genetically encoded GPCR-based fluorescent sensors showed promise to detect enkephalin^17^ and oxytocin^18^ *in vitro*, respectively. The “iTango” sensor allows single-cell resolution and nanomolar sensitivity for selected neuropeptides, such as vasopressin^19^ and oxytocin^20^, but provides poor temporal resolution (24-48 hours transcription). Other techniques such as neuropeptide release reporters detect real-time neuropeptide trafficking events but do not report information on specific neuropeptides^21^. Cell-based neurotransmitter fluorescent engineered reporters (CNiFERs), on the other hand, provide a potential solution to selectively detect nM concentrations of peptides *in vivo*. CNiFERs utilize a clonal HEK293 cell that is engineered to express a specific GPCR and amplifies the signal with a genetically-encoded fluorescence resonance energy transfer (FRET)-based Ca^2+^ sensor to detect neurotransmitters^22–24^. CNiFERs have been implanted in target brain regions to detect endogenous release of acetylcholine^22^, differentiate similar monoamines (dopamine vs. norepinephrine)^23^, have a spatial resolution defined by the microscale implants, and are fast enough to be implemented in a closed-loop reward feedback system^25^.

In this work, we developed an all-optical approach and demonstrate, for the first time, *in vivo* neuropeptide release and detection by photo-released neuropeptide from nanovesicles and CNiFERs sensor. The nanovesicles consist of a phospholipid liposome core and gold coating and can be remotely stimulated by near-infrared light to photo-release small volume (fL to pL) of biomolecules under tight temporal and spatial control. For detection, we developed a new cell-based neurotransmitter fluorescent engineered reporter (CNiFER) that is sensitive to nanomolar (nM) concentrations of somatostatin (SST), a 14-amino acid cyclopeptide expressed throughout the brain. Our approach takes advantage of the high spatial and temporal resolution of two-photon imaging system. The volume transmission of SST was measured in mouse cortex and we found the maximal diffusion distance to activate GPCR is 220 μm. We further demonstrate that the neuropeptide transmission and GPCR signaling is faster in hyaluronan-deficient mouse neocortex, a diseased condition indicated in neuroinflammation^26^ and various neurodegenerative diseases including Parkinson’s disease^27^.

## Results

### Tool development and characterization for neuropeptide release and sensing

Here, we focused on somatostatin-14 (SST), an endogenous cyclic 14 amino acid peptide hormone, because it is widely expressed in GABAergic cortical neurons but little is known about its function^28, 29^. SST was encapsulated in nanovesicles prepared by 1,2-dipalmitoyl-*sn*-glycero-3-phosphocholine (DPPC) and cholesterol (4:3 molar ratio) and photoreleased with femtosecond laser excitation (**Fig. 1a, Supplementary Fig. 1**). Gold-coating offers nanovesicles strong absorption in the near-infrared range (700-900 nm) and increases the hydrodynamic diameter to ~200 nm (**Fig. 1b, Supplementary Fig. 1**). Clusters of small gold nanoparticles were observed on the nanovesicle surface by transmission electron microscopy (TEM, **Fig. 1c**). The SST content in gold-coated nanovesicles (Au-nV-SST) remained over 75% after 24 hours of incubation with peptidase α-chymotrypsin, while the half-life of degradation for free SST was 16 mins (**Supplementary Fig. 1**). Au-nV-SST show good colloidal stability (UV-Vis spectra, particle size and polydispersity index) in 0.01M phosphate-buffered saline (PBS) at 4°C over two weeks (**Supplementary Fig. 1**). To characterize the SST release efficiency, Au-nV-SST flowing in a capillary was stimulated by femtosecond laser pulses and “tornado” scans using a multi-photon imaging system with two lasers tuned to different wavelengths, referred to as imaging laser and stimulation laser, respectively (**Supplementary Fig. 2**). Each tornado scan (200 μm, 35 ms) has a dwell time of 10 μs on a pixel, leading to 2.8 × 10^6^ pulses per scan (80 MHz laser repetition rate). The photoreleased SST concentration was measured by ELISA kits. **Fig. 1d** shows that the photorelease efficiency and the concentration of photoreleased SST increase with laser power and scan number. To analyze the photorelease in a small volume, we loaded gold-coated and calcein-containing nanovesicles (Au-nV-Cal) into 1 nL microwells (**Supplementary Fig. 3**); under these conditions calcein is encapsulated at a self-quenching concentration (75 mM) and the fluorescence increases upon release and dequenching. Calcein fluorescence increases significantly under stimulation laser pulses at 700-820 nm (100 mW power, 0.65 s duration), while showing no change or release under imaging laser pulses at 840-1000 nm, even at prolonged durations (15 mW power, 8 mins) (**Fig. 1e, Supplementary Fig. 3)**. The results suggest the stimulation laser leads to efficient molecule release, while the imaging laser does not photorelease encapsulated compounds and allows for long-term monitoring. Real-time fluorescent imaging shows that the average calcein fluorescence in a microwell (diameter: 60 μm) reached a maximal value at 0.15 s after a 65 ms tornado scan covering ~60 μm (**Fig. 1f, Supplementary Fig. 3)**. Thus, the photo-release from Au-nV-Cal has fast temporal resolution, completing within 0.15 s in the 60 μm tornado-scanned area.

**Fig. 1.**
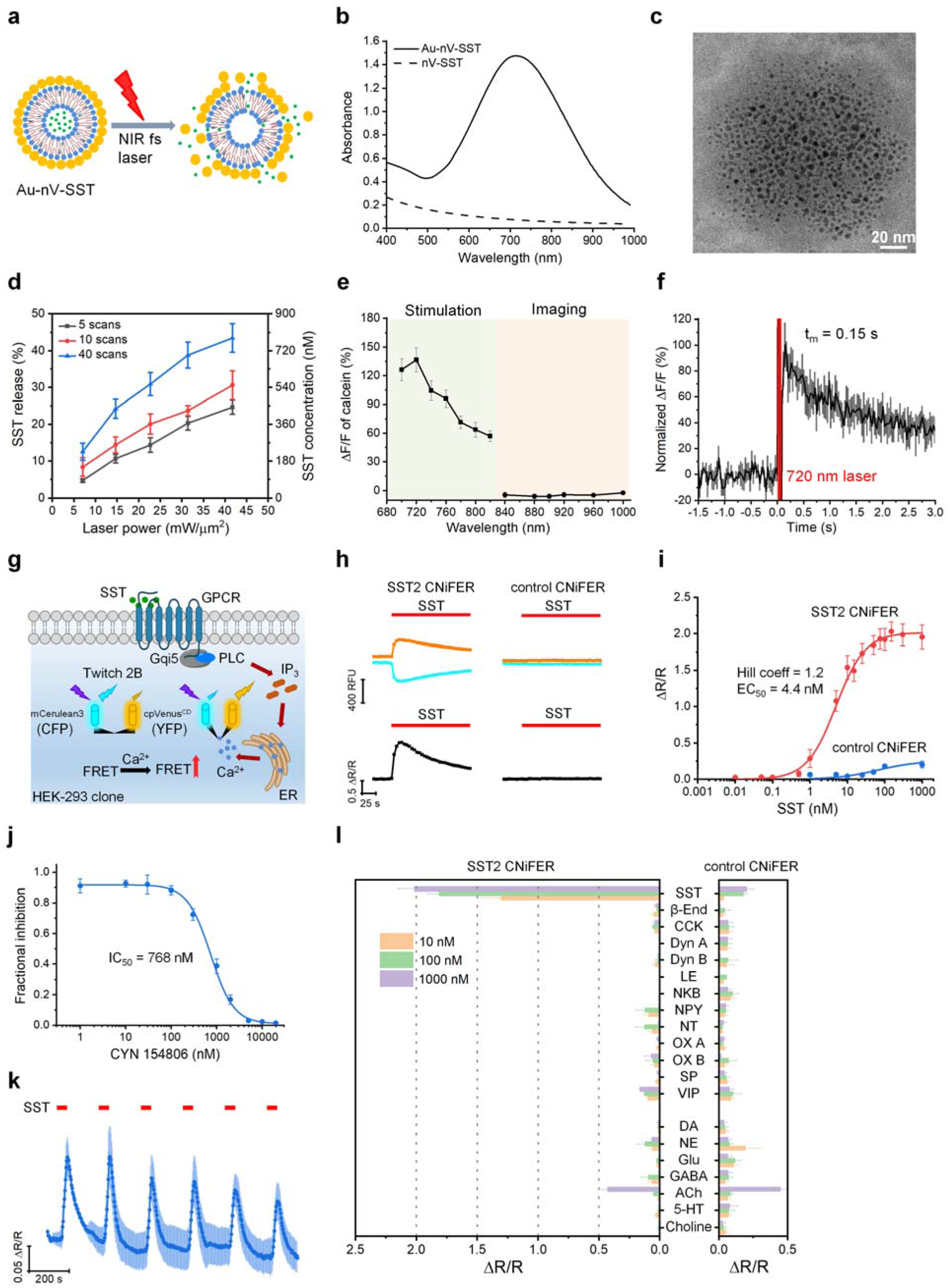
Development and characterization of neuropeptide photo-release and sensing techniques. **a**, Schematic of near-infrared (NIR) femtosecond (fs) laser pulses triggered release from gold-coated somatostatin-14 (SST) loaded liposomes (Au-nV-SST). **b**, UV-Vis spectra of Au-nV-SST and nV-SST. **c**, Transmission electron microscopy (TEM) image of Au-nV-SST. Scale bar: 20 nm. **d**, The efficiency and amount of SST released from nanovesicles under the irradiation of fs laser pulses (n = 3). **e**, Wavelength-dependent release of calcein under stimulation and imaging conditions (n = 5). Stimulation conditions: 100 mW, 10 tornado scans, 0.65 s, 700-820 nm. Imaging conditions: 15 mW, resonant scans, 8 minutes, 840-1000 nm. **f**, Release kinetics of calcein from nanovesicles stimulated at 720 nm for 65 ms (n = 5). **g**, Schematic of SST2 CNiFER signaling pathway. SST activates SST2 GPCR to induce Ca^2+^ cytoplasmic influx detected by the fluorescent genetically encoded Ca^2+^ detector (Twitch 2B). Twitch 2B contains mCerulean3 and cpVenus^CD^, which we refer to as CFP and YFP, respectively. **h**, FRET response from SST2 and control CNiFERs after application of 100 nM SST (red bar). Response in CFP (cyan) and YFP (yellow) fluorescence (top) leads to the change of FRET ration (ΔR/R, bottom). **i**, Dose-response curve for SST2 (n = 4) and control (n = 3) CNiFERs. Smooth curve shows best fit with Hill equation. **j**, Inhibition of SST (5 nM) activation of SST2 CNiFERs with different concentrations of the SST2 receptor antagonist (CYN 154806). **k**, FRET response for SST2 CNiFER with repeated 30 s applications of SST (5 nM, n = 4). **l**, FRET response with the indicated peptides and classical neurotransmitters (NTs) at three different concentrations for SST2 CNiFERs (n = 3-6) and control CNiFERs (n = 3-6). Data were expressed as Mean ± S.D.

To measure the release of SST *in vivo*, we engineered a new neuropeptide CNiFER that expresses the type 2 SST GPCR (SST2) along with a chimeric Gqi5 G protein and a high signal-to-noise FRET-based calcium sensor, Twitch 2B (**Fig. 1g**)^23, 30^. We first conducted a series of *in vitro* experiments to determine the sensitivity and specificity of the SST2 CNiFER. Application of SST to SST2 CNiFERs induces a FRET response, whereby the mCerulean3 (referred to as ‘CFP’) emission decreases and cpVenus^CD^ (referred to as ‘YFP’) emission increases with receptor activation (**Fig. 1h**). In contrast, SST application does not lead to a response in the control CNiFER. The SST2 CNiFER clone #5F3 was selected due to its high affinity binding for SST (EC_50_ = ~4 nM) and a large ΔR/R (~2), while the control CNiFER without the SST2 receptor showed little or no response with up to 1 μM SST (**Fig. 1i**). We further applied a selective SST2 receptor antagonist (CYN 154806) and observed complete inhibition of the SST-dependent CNiFER activation (**Fig. 1j**). To assess whether SST2 CNiFERs can respond to the repetitive release of SST, we measured the FRET response of SST2 CNiFERs to six pulses of SST. The amplitude of the FRET response (ΔR/R) decreased after the first pulse but appeared to reach a steady-state level of response with additional pulses of SST (**Fig. 1k**). Lastly, we screened the response of SST2 CNiFERs to a family of different neuropeptides and neurotransmitters at three different concentrations (10, 100, and 1000 nM), using a high-throughput fluorescent plate reader (**Fig. 1l**). The SST2 CNiFER shows a highly specific and robust response to SST. There is also a small response with acetylcholine at 1000 nM, but this occurs with both SST2 and control CNiFERs. Taken together, these results show that the SST2 CNiFER (#5F3) has nanomolar sensitivity, exhibits a large FRET ratio response, and is highly specific for SST detection.

### Real-time photo-release and monitoring of somatostatin-14

Next, we investigated the feasibility of simultaneously photoreleasing SST from nanovesicles and measuring SST with SST2 CNiFERs *in vitro* and *in vivo*. SST2 CNiFERs were grown in a monolayer and imaged with a two-photon microscope (**Fig. 2a**). Au-nV-SST (300 nM of SST) was applied to the CNiFERs *via* bath perfusion, and stimulated by 60 μm tornado scans at 720 nm (300 mW, 64 scans). Real-time two-photon fluorescent images show that the ratio of YFP and CFP of CNiFERs around the stimulation area dramatically increases after stimulation (**Fig. 2b**). The effect of photorelease is further visualized by the heat map of the FRET response (ΔR/R) for a total of 180 cells analyzed (**Fig. 2c**). Note that the cells far away have a delayed and smaller FRET response compared with cells closer to the tornado scan area (**Supplementary Fig. 4**). Furthermore, the FRET response from SST2 CNiFERs increases with the stimulation power and scan number. Importantly, there is no CNiFER response observed for photo-stimulation of gold-coated empty nanovesicles (Au-nV, therefore no SST release) or SST-loaded nanovesicles without gold coating (nV-SST) (**Supplementary Fig. 4**). These results suggest that the photoreleased SST can be detected by SST2 CNiFERs in an *in vitro* environment in real time.

**Fig. 2.**
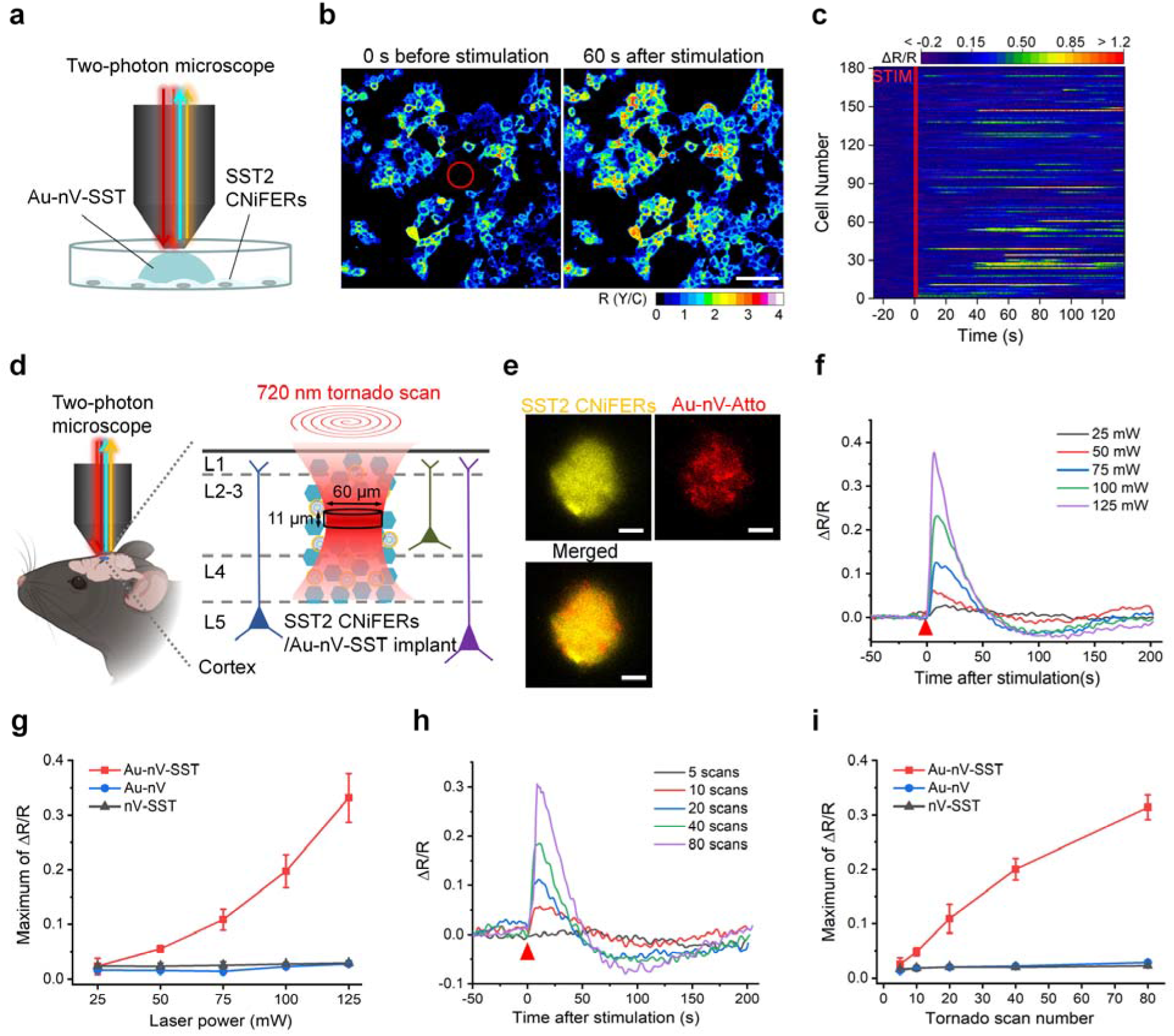
Real-time photo-release and monitoring of SST. **a**, Schematic of photo-releasing and monitoring SST on cultured SST2 CNiFERs by a two-photon microscope. **b**, Representative images of the ratio of CFP and YFP (R(Y/C)) of SST2 CNiFERs before and after laser stimulation on Au-nV-SST (720 nm, 300 mW, 64 scans). Tornado scans with a diameter of 60 μm was performed in the center (marked as red circle) at 0 s. Scale bar: 100 μm. **c**, Heat map of FRET response (ΔR/R) within individual cells. STIM represents the photo-stimulation at 0 s. **d**, Schematic of implantation of SST2 CNiFERs/Au-nV-SST mixture into mouse cortex. Au-nV-SST was stimulated by tornado scans (diameter: 60 μm) at 720 nm with an axial resolution of 11 μm. **e**, Representative two-photon fluorescent images of SST2 CNiFERs (E_x_: 900 nm; E_m_: 520-560 nm) and Atto 647N-labeled Au-nV (E_x_: 1100 nm; E_m_: 575-645 nm) at the depth of 200 μm in mouse cortex. Scale bar: 25 μm. **f**,**g**, FRET change (ΔR/R) trace (**f**) and maximum of ΔR/R (**g**) of the SST2 CNiFERs implant under 720 nm laser stimulation with different laser power (40 tornado scans, n = 3 implants from 3 mice per group). **h**,**i**, FRET change (ΔR/R) trace (**h**) and maximum of ΔR/R (**i**) of the SST2 CNiFERs implant under 720 nm laser stimulation with different scan number (100 mW, n = 3 implants from 3 mice per group). Au-nV: gold-coated empty nanovesicles. nV-SST: SST-loaded nanovesicles without gold coating. Data were expressed as Mean ± S.D.

To test the photorelease and monitoring of SST *in vivo*, we implanted the mixture of SST2 CNiFERs and fluorescent dye-labeled Au-nV (Au-nV-Atto, Atto refers to Atto 647N) into mouse somatosensory cortex and imaged within 2-4 hours under two-photon microscope with head-fixed mice (**Fig. 2d**). The YFP (520-560 nm) from CNiFERs and red fluorescence (575-645 nm) from Au-nV-Atto could be detected under imaging. **Fig. 2e** shows that the local distribution of Au-nV-Atto matched well with that of CNiFERs, suggesting the limited diffusion of nanovesicles in the brain (z-stack images in **Supplementary Fig. 5**). Upon 720 nm light stimulation and the subsequent SST release, we observed strong FRET increase that is dependent on the laser power and scan number (**Fig. 2f and 2h**). Photo-stimulation on Au-nV or nV-SST does not produce a FRET response in the SST2 CNiFERs implants even at highest laser power and scan numbers (125 mW and 80 scans), suggesting that the CNiFERs response is specific to the photoreleased SST (**Fig. 2g and 2i)**. Since the photorelease is a result of one-photon stimulation (laser stimulation matches absorption of nanovesicles), we further analyzed the photon transport by numerical simulation and obtained a stimulation volume of ~10 fL (0.93 μm × 0.93 μm × 11.3 μm) with a single pulse. A tornado scan of 60 μm diameter circle gives a stimulation volume of 32 pL (**Supplementary Fig. 6 and Table 1**). Stimulation of the 32 pL volume at 100 mW and 40 scans leads to an estimated photorelease of 1.2 × 10^8^ molecules, or 100 nM at the CNiFERs implant (**Supplementary Table 2**), a condition that may mimic volume transmission *in vivo*^31^. These results demonstrate physiological concentrations of SST can be photoreleased from nanovesicles and detected by SST2 CNiFERs *in vivo*.

### Measurement of somatostatin-14 volume transmission *in vivo*

Now, having a toolbox with both a photo-releasable SST and a SST detector, we designed an experiment to measure SST signaling over distance *in vivo* in real-time. We hypothesized that SST diffusion could be detected by implanting two distinct SST2 CNiFER clumps, and measuring the FRET response as SST diffuses from one clump to the other. To accomplish this, we implanted one cluster of SST2 CNiFERs co-mixed with Au-nV-SST (defined as the *core* implant) in the mouse cortex (**Fig. 3a**), and then implanted one or two additional SST2 CNiFER clumps at adjacent sites (defined as the *satellite* implant). **Fig. 3b** shows a core (ROI-1) and two satellite implants (ROI-2, ROI-3) at 200 μm depth in the mouse somatosensory cortex. Photostimulation of Au-nV-SST in the core implant (ROI-1) triggered SST release that was detected first by the core implant and then by satellite implants (**Fig. 3c**). The peak CNiFER responses at satellite implants (ROI-2 and ROI-3) were smaller and occurred after a time delay from the core implant (ROI-1), indicating the dilution and diffusion delay for the neuropeptide, respectively. For control, no response was observed when stimulating satellite implants that did not contain SST nanovesicles. We then systematically varied the distances between core and satellite SST2 CNiFER implants and determined the time delay over different distances (**Fig. 3d**). The time to activate SST2 CNiFERs positioned a short distance (less than 130 μm) was below 10 s and slowly increased with distance increasing, while it dramatically increased within larger distances (150-220 μm). Interestingly, no CNiFER response was measured at distances >220 μm, suggesting a maximum distance that the photoreleased neuropeptide is able to travel and produce a FRET response.

**Fig. 3.**
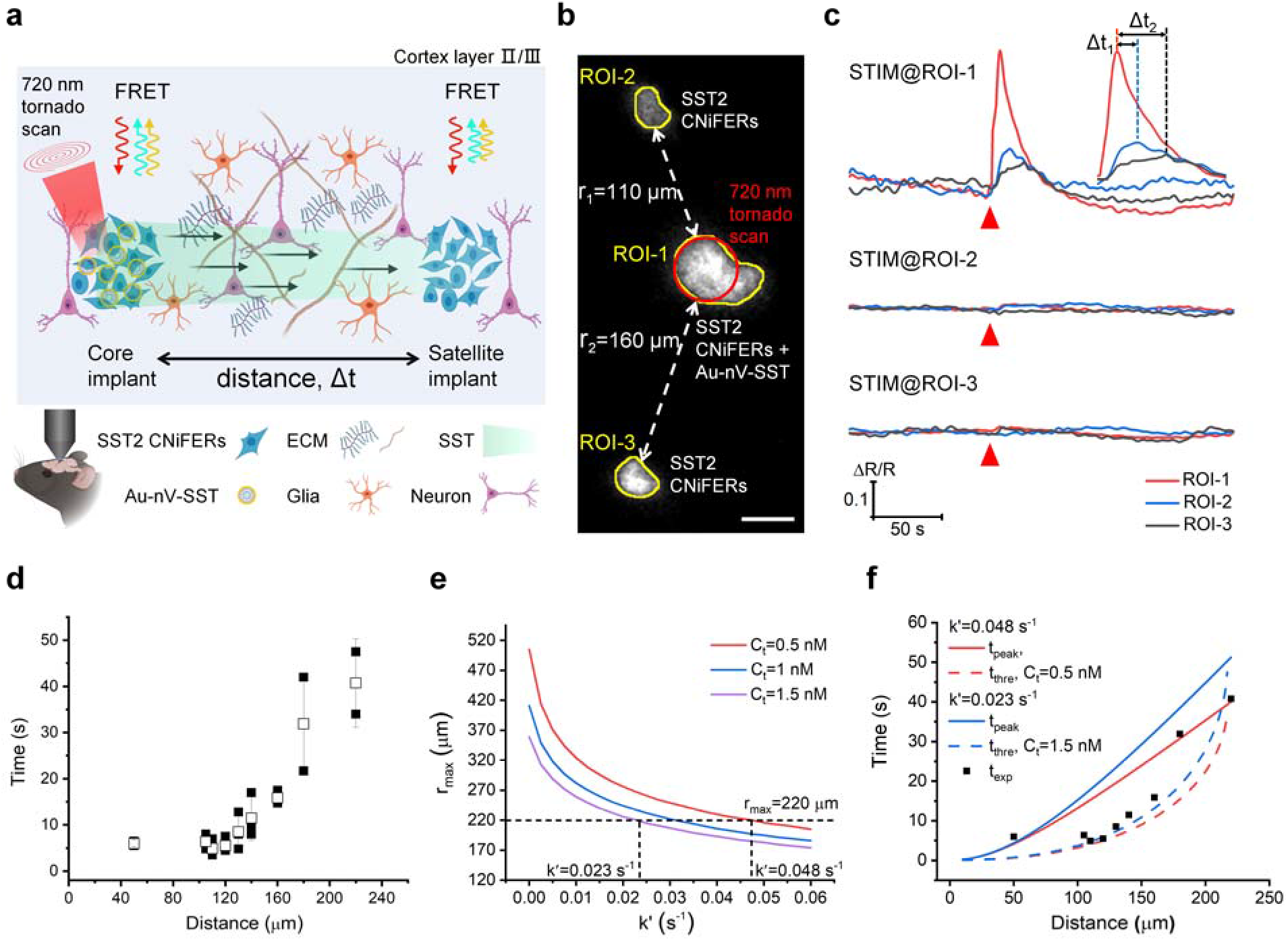
Measurement of SST volume transmission *in vivo*. **a**, Schematic of SST transmission measurement by implanting two clusters of SST2 CNiFERs, in which the core implant (left) is mixed with Au-nV-SST. **b**, Two-photon fluorescent image of SST2 CNiFERs at the depth of 200 μm in mouse cortex (E_x_: 900 nm; E_m_: 520-560 nm). The implant in the center (ROI-1) was mixed with Au-nV-SST, while the other two implants (ROI-2, ROI-3) were just SST2 CNiFERs alone. Scale bar: 50 μm. **c**, The response curves of SST2 CNiFERs when stimulating (STIM) at different regions (720 nm, 100 mW, 40 scans). The enlarged curve shows the time intervals between the peaks of response. **d**, Summary of SST transmission distance and corresponding time measured in normal mice brains (n = 9 mice, 2-4 repeated measurements for a pair of implants). Blank squares represent the mean values. **e**, Predicted maximal diffusion distance (*r_max_*) as a function of loss rate (k’) with total SST released number Q = 1.2 × 10^8^ and effective diffusion coefficient D* = 8.9 × 10^-7^ cm^2^•s^-1^ (from optical integrative imaging measurement). The horizontal dash line represents the experimental *r_max_* = 220 μm. **f**, The time (t_peak_) to reach maximal concentration and the time (t_thre_) to reach the threshold concentration as a function of distance. Black squares represent the mean values of measured diffusion time (t_exp_) at different distances. Data were expressed as Mean ± S.D.

We next compared our distance-dependent signaling measurements with a theoretical diffusion model. We first measured the effective diffusion coefficient (D*) of fluorescently labelled SST, fluorescein-5(6)-carbonyl-somatostatin-14 (5(6)-FAM-SST), by the integrative optical imaging (IOI) method in acute brain slices (**Supplementary Fig. 7,** D* = (8.9 ± 1.3) × 10^-7^ cm^2^•s^-1^)^32, 33^. Since the cyclic structure of SST remains the same in 5(6)-FAM-SST, this method gives a reasonable estimate of the effective diffusion coefficient (D*) for SST in brain tissue. We analyzed theoretical prediction and experimental measurement of the maximal signaling distance (*r_max_*), defined as the farthest location where the peak SST concentration reaches threshold for CNiFERs detection (C_t_). With the effective diffusion coefficient (D^*^), nanomolar sensitivity of CNiFERs (C_t_ = 0.5-1.5 nM, see Methods), and estimated number of photoreleased SST molecules (1.2 × 10^8^), the model gives a loss rate constant (k’) as a fitting parameter in the range of 0.023 s^-1^ - 0.048 s^-1^ (**Fig. 3e**). Here the loss rate constant k’ is a contribution of neuropeptide binding, uptake and degradation during diffusion in extracellular space (**Fig. 3a**). This estimate seems to be within the range of previous estimates with neuropeptide half-life ranging 1-3 minutes in plasma (for SST) ^34^, and ~20 mins in brain cerebrospinal fluid (CSF, oxytocin and vasopressin)^35^. With these transport parameters, we calculated the time for the released neuropeptide to diffuse and reach the threshold concentration (C_t_, Equation 2) and peak concentration (Equation 4) at different distances. The experimental data falls within the predicted times, which envelop the range of the time for the CNiFERs to sense and respond to the released neuropeptide (**Fig. 3f**).

### Somatostatin-14 volume transmission in hyaluronan-deficient mouse brains

To test these findings of SST diffusion, we applied the all-optical approach to probe SST volume transmission in mice with an altered extracellular environment. The geometry of brain extracellular space and the composition of extracellular matrix play import roles in controlling the local diffusion of endogenous signaling molecules or nutrients^6, 36^. Hyaluronic acid (HA) is one of the major components of brain extracellular space and forms high molecular weight long-chain molecules, a physical barrier for molecular diffusion. To alter volume transmission, we injected hyaluronidase (hyase) into the lateral ventricles of adult mouse brain and measured the SST diffusion two days after hyase injection. The reduction of HA was confirmed by a significant decrease in the hyaluronan binding protein (HABP) immunostaining (**Fig. 4a and 4b**). Quantitative analysis shows that the intensity of HABP decreased to 23% of that in normal brains (**Supplementary Fig. 8**). Importantly, no differences in cell nuclei number or size were observed in the same cortical area compared with the control group (**Supplementary Fig. 8**), indicating no significant cell loss.

**Fig. 4.**
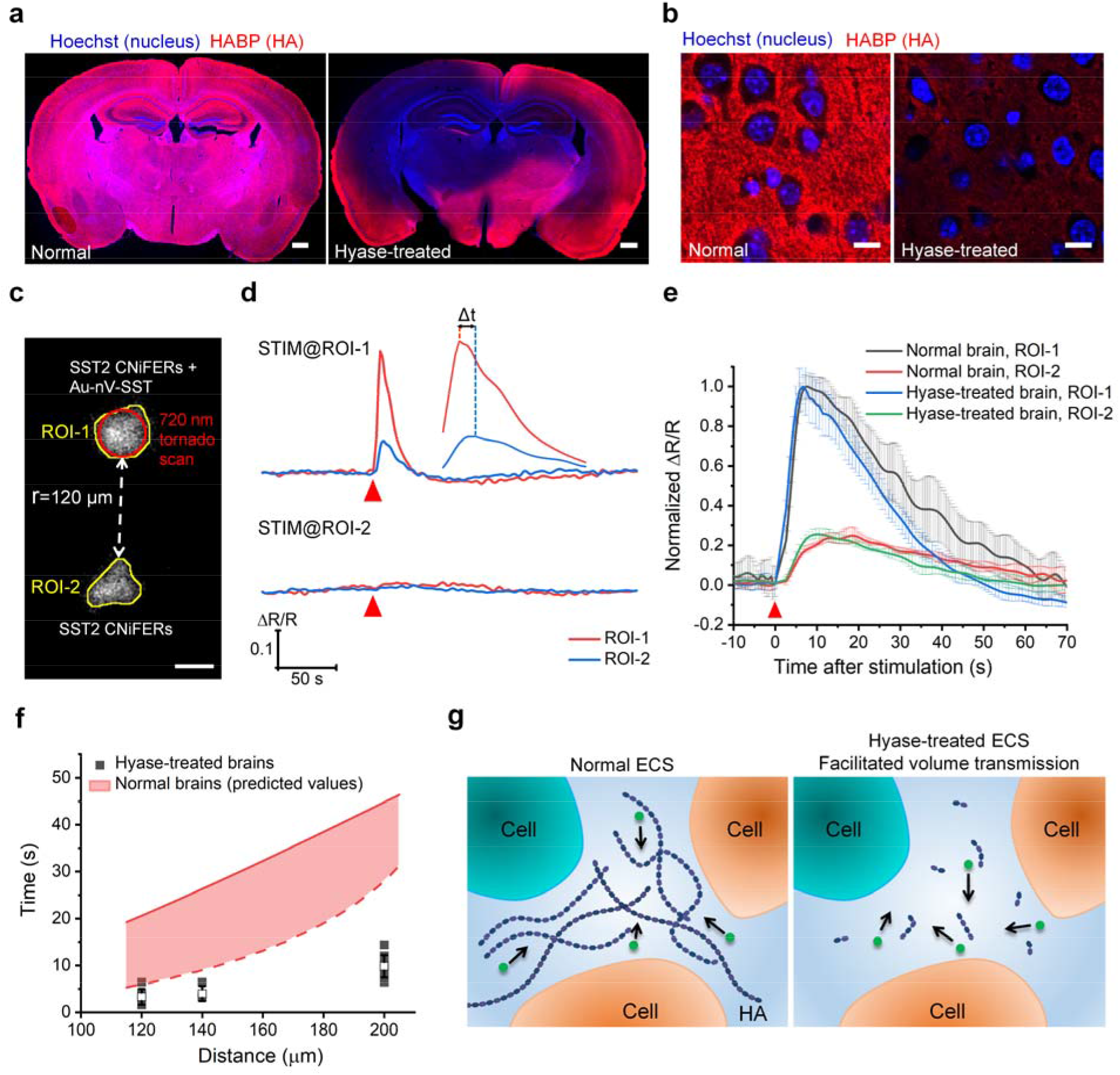
SST volume transmission in hyaluronan-deficient brains. **a**, Representative images of HABP-labeled (red) brain sections from normal mouse and hyaluronidase (hyase)-treated mouse. Nuclei were stained by Hoechst 33342 (blue). Hyase was injected into the left ventricle of mouse brain. Scale bar: 500 μm. **b**, Confocal images of brain cortex in the left hemisphere with high magnification. Scale bar: 10 μm. **c**, Two-photon fluorescent image of SST2 CNiFERs implanted at the depth of 200 μm in mouse cortex. The top implant (ROI-1) has SST2 CNiFERs mixed with Au-nV-SST, while the bottom implant (ROI-2) contains only SST2 CNiFERs. Scale bar: 50 μm. **d**, The response curves of SST2 CNiFERs when stimulating (STIM) at different regions (720 nm, 100 mW, 40 scans). The enlarged curve shows the time intervals between the peaks of response. **e,** Comparison of normalized ΔR/R traces of the core implants (ROI-1) and satellite implants (ROI-2) at the distance of 140 μm in normal and hyase-treated mouse brains (n = 3 measurements). **f**, Comparison of the time to activate GPCR at corresponding distance in hyase-treated mouse brains (n = 6 mice for total, 2-4 repeated measurements for a pair of implants) with the predicted time range in normal mouse brains (acquired from Fig. 3f). Red solid line shows the time to reach peak concentration at k’ = 0.023 s^-1^, while the red dash line shows the time to reach the threshold concentration (C_t_ = 1.5 nM and k’ = 0.023 s^-1^). **g**, Schematic of the change of brain extracellular space (ECS) and SST volume transmission under different conditions. Data were expressed as Mean ± S.D.

With the HA-deficient mouse brain model, we repeated the measurement of SST volume transmission using the core-satellite method as above (**Fig. 4c**). Stimulation of the core implant (ROI-1) containing both CNiFERs and Au-nV-SST led to a large increase in the CNiFER FRET response in the two implants, while no response was observed upon the stimulation of the satellite implant (ROI-2) containing only the CNiFERs (**Fig. 4d)**. Comparison of the core-satellite responses at the same distance (140 μm) suggests that the peak time has a faster rise and decay for satellite implant (ROI-2) in hyase-treated brain compared with normal brain (**Fig. 4e**). The CNiFER responses at three distances occurred much faster in the hyase-treated brain compared with normal brains (**Fig. 4f**). Thus, our results suggest that the HA reduction leads to a faster neuropeptide volume transmission in the HA-deficient mouse brains due to reduced hindrance from long-chained HA (**Fig. 4g**).

## Discussion

Current methods for precise temporal and spatial control of neuropeptide release and its detection *in vivo* in real-time are lacking in the field^4^. Here, we provide two new complementary neurotechniques to address this deficiency. The first technique provides the ability to package a neuropeptide in small gold-coated nanovesicles and photo-release in discrete regions in the brain. The second technique involves the creation of the SST2 CNiFER that detects nM concentrations of SST in extrasynaptic regions in the brain. With these tools, we systemically measured the time for SST signaling and transmission across a large range of distances and determined the maximum signaling distance with a bolus release of neuropeptides.

This all-optical approach represents a new platform to probe the extrasynaptic signaling through the extracellular space (ECS) in both healthy and diseased brains. The key feature of brain extracellular matrix (ECM) is the abundance of hyaluronic acid (HA) and HA-binding proteoglycans^37^, which are altered and can act as potential biomarker in many brain diseases including neuroinflammation^26^, Alzheimer’s disease (AD)^38^, Parkinson’s disease (PD)^27^, and tumor progression and metastasis^39^. Many methods have been developed to measure the diffusion coefficient of molecules in the brain ECS, such as integrative optical imaging (IOI)^32^ and real-time iontophoresis with ion-selective electrodes^40^, or super-resolution imaging of ECS structure recently^5, 7^. While these methods provide a useful tool to measure the diffusional and structural properties of ECS, few methods can probe the neuropeptide signaling through the ECS. Our approach provides an important new dimension to the increasingly relevant ECS and its role in extrasynaptic signaling and transmission by directly photoreleasing and sensing neuropeptides through GPCR. Specifically, this approach differentiates from the dye diffusion measurement and models the volume transmission better in that there are several steps between neuropeptide release and CNiFER response at a distance, including neuropeptide diffusion, binding to the GPCR, and subsequent signaling that lead to the measured Ca^2+^ response. Therefore, it is complementary to the existing methods and provide a new perspective to study the volume transmission. Furthermore, by combining with IOI which measures the effective diffusion coefficient (D*) of 5(6)-FAM-SST, we estimated the *in vivo* loss rate constant for SST (k’) to be in the range of 0.023 - 0.048 s^-1^, a measurement that is not possible by each method alone.

One important consideration of the all-optical approach is the impact of local brain heating on the measurement. Tissue heating is a common concern for multiphoton techniques for *in vivo* imaging^41, 42^. Here the gold-coating absorbs single photon at 720 nm to trigger the molecular release from nanovesicles. To measure the tissue heating, we inserted a thermocouple probe in the brain during laser stimulation and imaging (**Supplementary Fig. 9**). Our results suggest that the heating was predominantly from tissue absorption of the near infrared light (3 °C/100 mW), consistent with previous reports^42^. Importantly, the presence of gold-coated nanovesicles does not lead to additional temperature increase, suggesting the dominant absorption from the tissue. Furthermore, there are thermo-sensitive TRP channels on the membrane of SST2 CNiFERs, and our experimental results show no observable response to the stimulation alone without SST release (**Fig. 2g and 2i**). Therefore, the tissue heating is very limited and does not have a significant impact on the signaling measurement reported here.

The technologies presented here represent a significant advance over current methodologies. The *in vivo* optical stimulation provides a high spatial (μm, or fL volume) and temporal (subsecond) resolution to control neuromodulators release compared with other approaches using magnetic^43^ or acoustic methods^44, 45^. The *in vivo* neuropeptide detection by CNiFERs measures nanomolar neuropeptide concentration that is physiologically relevant, highly specific, and in real time. By contrast, microdialysis is offline measurement^46^ and fast scanning cyclic voltammetry may not differentiate peptides with clear oxidation signals on carbon fiber probes^47^. A rough estimation suggests that a neuron releases 10^6^ peptide molecules per second upon stimulation, and possibly higher in the hypothalamic neurons (10^7^ molecules per second)^31^. Therefore, the estimated neuropeptide release (10^8^) is close to neuropeptides released from a local region of neurons (10 ~ 100). Our work mainly focuses on the molecular diffusion in layers II/III of mouse cortex, while diffusion is not only heterogeneous but also anisotropic in brain regions such as corpus callosum, hippocampus, and striatum^6^. It will be important to compare our measurements with volume transmission in different brain regions in the future.

## Methods

### Materials

1,2-Dipalmitoyl-*sn*-glycero-3-phosphocholine (DPPC, 63-89-8, >99%) and cholesterol (ovine wool, 57-88-5, >98%) were purchased from Avanti Polar Lipids, Inc. Calcein sodium salt (644.5 g/mol, 108750-13-6) was purchased from Alfa Aesar. Somatostatin-14 (SST, 38916-34-6, 1637.9 g/mol, >99%) and Fluorescein-5(6)-carbonyl-somatostatin-14 (5(6)-FAM-SST, 1996.2 g/mol, >96%) were purchased from Bachem, Inc. Tetrachloroauric(III) trihydrate (HAuCl_4_·3H_2_O, 16961-25-4, 99.9%), *L*-ascorbic acid (50-81-7, 99%) and Atto 647N-labeled 1,2-dipalmitoyl-*sn*-glycero-3-phosphoethanolamine (Atto 647N DPPE, ≥90.0%) were purchased from Sigma-Aldrich. Gibco™ Dulbecco’s Modified Eagle’s Medium (DMEM, high glucose) with GlutaMAX™ supplement, trypsin–ethylenediamine tetra-acetic acid, fetal bovine serum and penicillin-streptomycin were purchased from ThermoFisher Scientific. All other chemicals were analytical grade.

### Preparation and characterization of gold-coated liposomes (Au-nV)

Gold-coated liposomes were prepared by the thin film-rehydration method as reported before with minor modifications^15, 48^. First, lipid film was prepared after dissolving 10 mg of DPPC and 4 mg of cholesterol in 0.5 mL of CHCl3. 0.5% of Atto 647N-DPPE was added as fluorescent label. Then the film was dried under vacuum and subsequently hydrated with 10 mM phosphate buffered saline (PBS) containing either calcein (75 mM) or somatostatin-14 (SST, 1.22 mM) under 55 °C for 1 h, followed by 5 freeze-thaw cycles (1 minute in liquid N_2_ and 2 minutes in 55 °C water bath, respectively). Afterwards the suspension was extruded through 200 and 100 nm polycarbonate membranes (Whatman, USA) for 21 passages each using a Mini Extruder (Avanti Polar Lipids, USA). Free calcein or SST was removed by size-exclusion chromatography with Sephacryl^®^ S-1000 column (Cat. # 17-0476-01, GE Healthcare). Second, gold nanoparticles were decorated onto the liposome surface *via* the reduction of gold chloride (10 mM) with ascorbic acid (40 mM) following similar procedures as reported^15, 48^.

The hydrodynamic sizes of nanovesicles were determined by dynamic light scattering measurement (Malvern Zetasizer Nano ZS). UV-Vis absorption spectra of nanovesicles in PBS were obtained with a plate reader (Synergy 2, Bio-Tek). The morphology of nanovesicles was observed by a transmission electron microscope (TEM, JEOL-1400+) at an accelerating voltage of 150 keV. Liquid chromatography-mass spectrometry (LC-MS/MS) was used to explore the kinetics of SST degradation by peptidase α-chymotrypsin (EC3.4.21.1, Cat. # C4129, Sigma-Aldrich). The mass ratio of SST to α-chymotrypsin was 100:1 and 50% methanol was used to lysis liposome right before measurements. Gradient mixture of acetonitrile and water from 10% to 100% was used as mobile phase in Agilent 1100 HPLC and SST was quantified by mass spectrometry (Sciex QTRAP 4000) with an internal standard ([ring-D_5_]Phe^6^)-somatostatin-14 (Cat. # 4072026, Bachem).

### *In vitro* photorelease efficiency

Calcein release from gold-coated liposomes (Au-nV-Cal) was investigated by loading the sample into a microwell array with 20,000 microwells and 60 μm diameter for each microwell (QuantStudio™ 3D digital PCR chip). This allows imaging the calcein release and fluorescent intensity increase in a small nL volume without diluting the fluorescence to background levels. Real-time imaging and photo-stimulation of nanovesicles were conducted on a two-photon microscope as detailed below. Calcein release at different stimulation wavelengths, power, and duration was characterized by the maximum fluorescence intensity increase (F_m_) in a well after stimulation relative to the baseline: ΔF/F = (F_m_-F_0_)/F_0_.

A capillary flow system was used to test the release efficiency of SST. Briefly, SST-loaded gold-coated liposomes (Au-nV-SST) were flowed through a square glass capillary with inner diameter of 200 μm. Tornado scans (diameter: 200 μm; constant linear velocity) of femtosecond laser pulses (MaiTai HP DeepSee-OL, Spectra-Physics, 80 MHz, 100 fs pulse width) were conducted on the capillary with a 5X objective (MPLN5X, Olympus, NA 0.1). The flow rate was set at 2, 1 and 0.25 μL/min by a syringe pump and gives the nanovesicle exposure at 5, 10, 40 scans, respectively. Each pixel scan lasts 10 μs and corresponds to 800 pulses. Laser pulses at 720 nm with different power (75, 155, 245, 340, 450 mW, measured after the objective) were tested. Liposomal suspension after irradiation was collected at the end of the capillary. SST concentration was measured by an enzyme immunoassay kit (Cat. # S-1179, Peninsula Laboratories International, Inc) according to the provided protocol.

### Construction of SST2 CNiFER

The CNiFER system is a modular design that can be applied to any identified GPCR. HEK293 cells were obtained from the ATCC. Based on our previous CNiFERs^23^, we increased the brightness and sensitivity by replacing the TN-XXL FRET based Ca^2+^ sensor with Twitch 2B (Tw2B)^30^. Tw2B contains mCerulean3 and cpVenus^CD^, which we refer to as CFP and YFP, respectively, for simplicity. Tw2B has a ΔR/R of 800% and a K_d_ of 200 nM for Ca^2+^. Because the SST2 receptor couples to Gi/o G proteins, we also co-expressed a chimeric G Protein Gqi5 that redirects Gi/o receptors to the Gq signaling pathway^23^. With expression of Gqi5 and Tw2B in HEK293 cells, we detected a significant FRET response with ACh, and subsequently performed CRISPR to knockout the endogenous human muscarinic M4 receptor. This modified HEK293 cell, clone #128C3 (Tw2B, Gqi5, hM4-CRISPR), had a significantly smaller ACh response and was used to generate the SST2 CNiFER, as described previously^23^. Briefly, hSSTR2 cDNA was cloned into pCDH vector (pCDH-CMV-MCS-EF1-Puro) to produce a Lentivirus that transduced 128C3 HEK293 cells. Fluorescence-activated cell sorting (FACS) was performed following puromycin selection, and a single clone (#5F3) was isolated that exhibited high sensitivity and ΔR/R with SST. The control CNiFER is clone 128C3, which lacks the SST2 receptor. CNiFERs were cultured in DMEM medium supplemented with 10% fetal bovine serum.

### Pharmacological evaluation of CNiFERs

All agonist and antagonist experiments were carried out in a 96-well plate and a plate reader (Molecular Devices Flexstation III). CNiFERs were plated in a 96-well black bottom plate the day before the experiment. The drug plate contained concentrated agonists or antagonists dissolved in artificial cerebrospinal fluid (ACSF) (125 mM NaCl, 5 mM KCl, 10 mM D-Glucose, 10 mM HEPES-Na, 3.1 mM CaCl_2_, 1.3 mM MgCl_2_). FRET was measured using 436 nm excitation and 485 nm (CFP) and 527 nm emission (YFP) every 4 sec for a total of 180 s. Agonist was added to the well after 30s and the peak FRET ratio was calculated for each well, averaged over triplicates, and then averaged over multiple runs. The dose-response curve was fitted with the Hill equation for activation (EC_50_), or the competitive inhibition curve was fitted with the Hill equation for inhibition (IC_50_).

To assess potential desensitization of the SST2 CNiFER, we examined the CNiFER response with multiple pulses of SST under fast perfusion. On Day 1, the CNiFERs were seeded on PDL-coated cover glasses (Diameter: 12mm; Thickness: #1) and the FRET response was measured on Day 2 with an inverted microscope (Nikon TE300) fitted with a 430nm LED (Prizmatix) for excitation, a dual bandpass dichroic (Chroma 51017bs), a filter wheel (Sutter Lambda) with CFP (Chroma ET485/30m) and YFP (Chroma ET535/50m) emission filters, and a CMOS camera (Andor Zyla). Imaging setup components were controlled by Micromanager software. Alternating CFP and YFP images were collected every 3 s. Six pulses of SST (5 nM, 30 s) in ACSF were delivered every 3 min. The solutions were delivered *via* a gravity-fed solenoid-controlled valve bank and 6-channel manifold (Warner). CFP and YFP fluorescence were measured in selected CNiFER ROIs and the ΔR/R calculated for each frame.

To monitor the SST release from Au-nV-SST *in vitro*, SST2 CNiFERs were seeded and cultured in 25 mm glass bottom dishes for 24 h. The cells were then washed with PBS and imaged by the two-photon microscope mentioned below. 250 μL of fresh ACSF and 50 μL of Au-nV-SST were mixed and added between the cells and the objective (25X, NA 1.05). After 10 minutes, photo-stimulation at 720 nm was performed besides the cells and the fluorescence of CNiFER was recorded at 0.5 s per frame.

### Animal preparation

Male C57BL/6 mice (two months old) were purchased from Charles River Laboratories, Inc. and were given free access to food and water before all experiments. Animal protocols were in accordance with NIH guidelines and approved by the Institutional Animal Care and Use Committee of University of Texas at Dallas and SUNY Downstate Health Sciences University. For surgery, mice (2-3 months old) were anesthetized with 1.5-2% isoflurane (Vedco, Inc) and fixed in a stereotaxic frame with ear bars. An open-skull cranial window with size around 3 mm x 3 mm was prepared by a drill in the somatosensory cortex for all the experiments. The mouse body temperature was maintained at 37 °C using a heat pad regulated by a rectal probe. Subcutaneous injections of 5% (w/v) glucose in saline were given every 2 h for rehydration. Dexamethasone (Sigma-Aldrich) was administered subcutaneously at the dosage of 5 mg/kg to prevent swelling of the brain and/or an inflammatory response. Buprenorphine SR-Lab (ZooPharm) was administered subcutaneously at the dosage of 1 mg/kg for post-operative analgesia. After nanovesicles or CNiFERs implantation, the mice were immobilized on a custom-built imaging frame and taken for two-photon imaging immediately. The mice were anesthetized with 1.5% isoflurane (Vedco, Inc) during imaging.

For hyaluronan-deficient mouse model, approximately 4 μL of 20 mg/mL hyaluronidase from bovine testes in PBS (Cat. # H3506, Sigma Aldrich) was injected intraventricularly at the coordinate of −0.5 mm A/P, +1.0 mm M/L and a depth of 2 mm two days before the diffusion measurement.

### CNiFERs/Au-nV-SST implantation in mouse cortex

SST2 CNiFERs were harvested from culture flask, centrifuged, resuspended in ACSF and filtered through a Cell Strainer (Corning^™^) with a pore size of 40 μm. The cells were then spined down and mixed with Au-nV-SST at a ratio of 10^6^ cells/μL. The SST2 CNiFERs/Au-nV-SST mixture (core implant) was loaded into a glass pipette with a tip diameter of 40 μm and 90-140 nL of mixture was injected into the neocortex 400-500 μm below the cortical surface within the following stereotaxic coordinates: −1.5 to 2.5 mm A/P, +2 to +3 mm M/L, relative to Bregma. Several clusters of SST2 CNiFERs without nanovesicles (satellite implants) were implanted into adjacent sites. A series of distances between core and satellite SST2 CNiFER implants were carried out to measure SST volume transmission.

### Two-photon imaging and stimulation

Au-nV and CNiFER cells were imaged with a multi-photon laser scanning microscope (FVMPE-RS, Olympus). FVMPE-RS was accompanied with stimulation laser (MaiTai HP DeepSee-OL, Spectra-Physics, 100 fs pulse width) and main scanner laser (Insight DS+ −OL, Spectra Physics, 120 fs pulse width) with a repetition rate of 80 MHz. Real-time imaging of SST2 CNiFER cells was realized by detecting the fluorescence of the genetically-encoded FRET-based Ca^2+^ sensor, Twitch 2B. The fluorescence emission in two channels (CFP 460-500 nm and YFP 520-560 nm) was collected by the 25x objective (XLPLN25XWMP2, Olympus, NA 1.05) when excited at 900 nm (less than 15 mW after objective). Atto 647N-labeled Au-nV (Au-Atto-nV) was detected by recording the fluorescence emission at 575-645 nm when excited at 1100 nm. Photo-stimulation on the nanovesicles was performed at 720 nm with a series of laser power (25, 50, 75, 100, 125 mW) and tornado scans (60 μm, 5, 10, 20, 40, 80 scans) to characterize the power- and scans number-dependent release of SST. Here tornado scan refers to scan setting for fast photostimulation and allows rapid scanning of a circular area. To measure the transmission of SST, 40 tornado scans (duration: 2.6 s) at 100 mW were stimulated on the core SST2 CNiFERs implants while monitoring the FRET changes on both the core and satellite implants at 0.8 s per frame (typically 400 frames).

### Temperature measurement during two-photon stimulation

The temperature change in the mouse brain during stimulation was monitored by implanting an IT-24P ultrafine flexible thermocouple probe (Physitemp) besides the core implant (SST2 CNiFERs/Au-nV-SST). The tip diameter of the probe was around 210μm and coated with CdSe quantum dots (Cat. # CSE600-10, NN-Labs) for visualization under two-photon microscope (E_x_: 900 nm, E_m_: 575-645 nm). The real-time temperature was recorded at 10 counts/s by an input module and customized Labview software. 40 tornado scans were performed at the core implant and adjacent blank sites at different power levels (50, 75, 100, 150, 200 mW after objective).

### Immunohistochemistry staining

In order to examine the changes of the extracellular matrix before and after hyaluronidase treatment, we used hyaluronic acid binding protein (HABP) to specifically detect the hyaluronan. Mice were transcranially perfused with 10 mM PBS (pH 7.4) followed by 4% paraformaldehyde (PFA) in PBS. Brains were extracted, postfixed overnight in 4% PFA, and dehydrated in 10% and then 30% (w/v) sucrose PBS solution at 4 °C for two days, respectively. Free-floating coronal sections of 30 μm thickness were cut using a cryostat (ThermoFisher). Before antibody incubation, the slices were incubated with 1% Triton X-100 and 4% bovine serum albumin (BSA) in PBS as the block buffer for 2 h under gentle shaking at room temperature. After washed three times with PBS, slices were incubated with biotinylated HABP (Amsbio # AMS.HKD-BC41, 1:200 dilution) for 48 h at 4 °C in a 0.2% Triton X-100 and 2% BSA PBS solution. Slices were subsequently washed three times with PBS, and incubated with streptavidin-Cy3 (ThermoFisher Scientific, Cat. # 434315, 1:1000 dilution) in 0.2% Triton X-100 and 2% BSA PBS solution, along with Hoechst 33342 (ThermoFisher Scientific, Cat. # 62249, 1:2000 dilution) to label nuclei for 2 h at room temperature under gentle shaking. After another PBS wash for 3 times, the slices were mounted with Fluoromount-G (SouthernBiotech). Imaging was performed with Olympus VS120 100-Slide scanning system using a 10X objective or Olympus FV3000RS laser scanning confocal microscope using a 40X objective.

### Brain slice preparation

Acute brain slices from C57BL6 mice (2 - 3.5 months old, n = 5) were prepared to measure the diffusion for fluorescently-labelled SST (5(6)-FAM-SST). Animals were first anesthetized with urethane (1.8 g/kg i.p.), and then were decapitated to extract the brain. A hemisphere of the brain was submerged in artificial cerebrospinal fluid (ACSF, containing 124 mM NaCl, 5 mM KCl, 26 mM NaHCO_3_, 1.25 mM NaH_2_PO_4_, 10 mM D-Glucose, 1.3 mM MgCl_2_ and 1.5 mM CaCl_2_, continuously bubbling with a mixture of 95% O_2_ and 5% CO_2_) precooled to 4 °C. Then, 400 μm thick coronal slices were prepared using a vibrating microtome. The slices were then placed on a net submerged in ACSF at room temperature for an hour, while the ACSF was continuously bubbled with a mixture of 95% O_2_ and 5% CO_2_. Slice containing somatosensory cortex was then transferred onto the imaging setup for experiment. The slice was placed in a submersion slice chamber and a peristaltic pump was used to perfuse the slice with ACSF (continuously bubbled with 95% O_2_ and 5% CO_2_ mixture) flowing at a rate of 2 mL/min. The temperature of the flowing ACSF was maintained at 33 ± 1 °C using an inline heater.

### Integrative Optical Imaging (IOI)

Fluorescein-5(6) was used to label SST at N-terminal of peptide sequence to produce 5(6)-FAM-SST with the molecular weight increased by 22%, while the cyclic structure of peptide remained the same. IOI technique was used to quantify diffusion coefficient of 5(6)-FAM-SST in a dilute 0.3% agarose gel and in the extracellular space of the somatosensory cortex region of acute mouse brain slices, respectively. IOI technique is described in detail previously^32, 33^. Briefly, a 1.5 mM solution of 5(6)-FAM-SST (in PBS) was pressure-injected from a single barrel glass micropipette into the brain tissue or dilute agarose gel (0.3% w/v in PBS; NuSieve GTG agarose, Lonza, Cat. # 50081) and a sequence of images were taken using a charged-coupled device (CCD) camera attached to an epifluorescence microscope. The steps of pressure injection and imaging were controlled automatically using a customized MATLAB-based program (IOIGUY). The distribution of the fluorescence signal in the images was then fitted to the diffusion equation using another customized MATLAB-based program (IDA). Nelder-Mead simplex algorithm was used for fitting (details about fitting explained in the reference^32^). Free diffusion coefficient in dilute agarose gel and effective diffusion coefficient in brain slices were acquired from this fitting. Since high concentration (mM) of 5(6)-FAM-SST was injected to the brain slices, the effect of peptide loss (k’) on this diffusion measurement is negligible.

### Data analysis

To analyze the CNiFERs response, ROIs were drawn by the segmentation plugins in Image J. The ratio of average fluorescence intensities in the two channels was calculated. Responses were quantified as the fractional change in the FRET ratio (ΔR/R), which was calculated by the following equation:

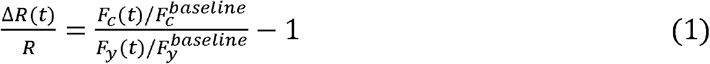

where *F_c_*(*t*) and *F_y_*(*t*) represent real-time fluorescence intensity in CFP and YFP channels, respectively; 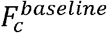 and 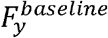 represent average fluorescence intensity before stimulation. Maximal responses were measured at the peak of ΔR/R after low-pass filtering (passband frequency: 0.1 Hz). The significant response was considered when maximal of ΔR/R ≥ 3δ (δ: Mean + S.D. of baseline).

### Modeling for the two-photon laser beam propagation

First, the distribution of light intensity in brain under the stimulation of femtosecond laser scans was obtained by a numerical model based on the beam propagation method (BPM) developed by Cheng et al^49^. The key idea of the BPM is to treat the scattering material as a series of planes that orthogonal to the initial light propagation direction. At each plane, we simulate the scattering by multiplying the local wavefront by a random phase term^49, 50^. Here we adopt a 2D-cylindrical BPM model with laser propagated along the z axis to predict the wavefront in the water and brain (**Supplementary Fig. 2 and Fig. 6**). The optical properties of brain cortex were extracted from the reference (**Supplementary Table 1**)^51^. The effective stimulation volume was calculated based on a cylinder model and setting full width of half maximum (FWHM) as an intensity threshold.

### Estimation of number of SST release (Q) *in vivo*

To estimate the amount of photoreleased SST *in vivo* (**Supplementary Table 2**), we first measured the SST concentration in the Au-nV-SST solution and after mixing it with the CNiFERs before implantation (C_sst_’ = 0.74 μM). Next, we measured the size of CNiFERs implant *in vivo* and approximated it as a cylinder with 80 μm in diameter and 400 μm height (**Supplementary Fig. 5**), giving a 2 nL in volume. This is much smaller than injected volume (average 120 nL) and we attribute to the water diffusion into the surrounding extracellular matrix. Since our imaging suggests limited nanovesicle diffusion (**Fig. 2e**), we argue that the nanovesicles remain in the CNiFERs implant. Lastly, we estimated the amount of photoreleased SST based on the release efficiency calibration curve. The effective stimulation volume (32 pL) and laser power density (3.6 mW/μm^2^) were acquired from the beam propagation model discussed above (**Supplementary Fig. 2 and Fig. 6)**. Based on the laser power intensity and calibration curve, we obtained the release efficiency (α = 13.8%) *in vivo* (**Supplementary Table 1)** and the amount of photoreleased SST (1.2 × 10^8^, **Supplementary Table 2**).

### Theoretical diffusion modeling

The diffusion equation is solved by taking a point-source paradigm in both space and time^6, 52^. The resulting concentration *C(r,t)* with respect to location *r* (distance from the source) and time *t* in the brain is listed here:

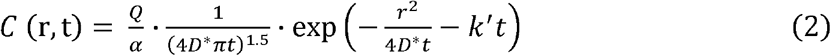

where *Q* is the total molecule number from the source; α is the volume fraction of brain extracellular space, which is set as 0.23 according to reference^6^; *D*^*^ is the effective diffusion coefficient *in vivo;* k’ is loss rate constant (binding, uptake, degradation etc.).

For a given distance (*r*), *C(r,t)* increases to reach the maximum concentration (*C*_peak_) for this location before decreasing with time. The time (*t_peak_*) to reach the maximum concentration for a given location *r*, can be calculated by solving the derivative of equation (2) while holding *r* as a constant:

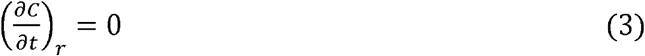

For SST diffusing in mouse cortex, the binding/uptake can be significant. Solving equation (3) results in the following function:

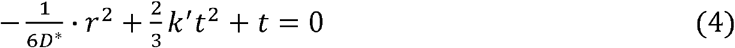

where D^*^ can be acquired from optical integrative imaging measurement. *t_peak_* will be calculated from equation (4) once the range of k’ is determined.

### Analysis of maximal diffusion distance

Maximal diffusion distance (*r_max_*) is defined as the farthest location where the peak SST concentration reaches threshold of detection (*C_t_*). The SST detection of limit by SST2 CNiFERs *in vitro* was calculated at 0.5 nM from the response curve in **Fig. 1i**. The *C_t_ in vivo* was then inferred in the range of 0.5-1.5 nM based on previous work^23^. *r_max_* was computed according to equation (2) by using a range of k’ (0 ~ 0.06 s^-1^) and *C_t_* (0.5, 1, 1.5 nM), given estimated number of released SST (*Q* = 1.2 × 10^8^) and *D*^*^ = 8.9 × 10^-7^ cm^2^•s^-1^.

### Statistical analysis

All data were collected in more than triplicate and reported as mean and standard deviation. Two-sample Student’s t-test in Origin 2021 software was conducted to determine p values and statistical significance. Single asterisk (*) indicates p < 0.05, double asterisks (**) indicate p < 0.01, and triple asterisks (***) indicate p < 0.001 as a measure of significance.

## Code availability

The custom MATLAB codes for theoretical diffusion modeling and data analysis used in this work are available upon request to Z.Q.

## Acknowledgements

We thank Dr. Oliver Griesbeck for providing the Twitch2B cDNA clone and discussions. This work was partially supported by National Science Foundation under award number 1631910 (Z.Q.), National Institute of Neurological Disorders and Stroke of the National Institutes of Health (NIH) under award number RF1NS110499 (Z.Q., P.A.S.), National Institute of Mental Health of NIH under award number R01MH111499 (P.A.S.), Human Frontier Science Program (HFSP) under award RGP0036/2020 (S.H.), and a postdoc research grant from the Phospholipid Research Center (Heidelberg, Germany) to H.X. Some schematics were made in part using BioRender.

## Author contributions

H.X., E.L., P.A.S. and Z.Q. conceived and designed the experiments. E.L., T.K. E.A. and Daniel Kircher fabricated and characterized the SST2 CNiFERs. E.L., C.M. and David Kleinfeld assisted in the microinjection of CNiFERs. H.O. performed the theoretical modeling for molecular diffusion. A.N. executed the diffusion measurement by the method of optical integrative imaging and analyzed the data under the supervision of S.H.. X.X. conducted performed the immunohistochemistry staining and assisted in data analysis. C.X. performed the theoretical modeling for two-photon laser beam propagation. J.Y. performed LC-MS/MS to measure the SST concentration. K.K. assisted in the animal experiments. X.L. and J.A.Z initiated the idea of the plasmonic nanovesicles. P.A.S. and David Kleinfeld initiated the construction of CNiFERs. H.X. performed the experiments, analyzed the results, wrote and revised the manuscript. P.A.S. and Z.Q. analyzed the results and revised the manuscript. P.A.S. and Z.Q. supervised the project. All authors discussed the results and commented on the manuscript.

