## Supplementary data for "Probing neuropeptide volume transmission in vivo by a novel all-optical approach"

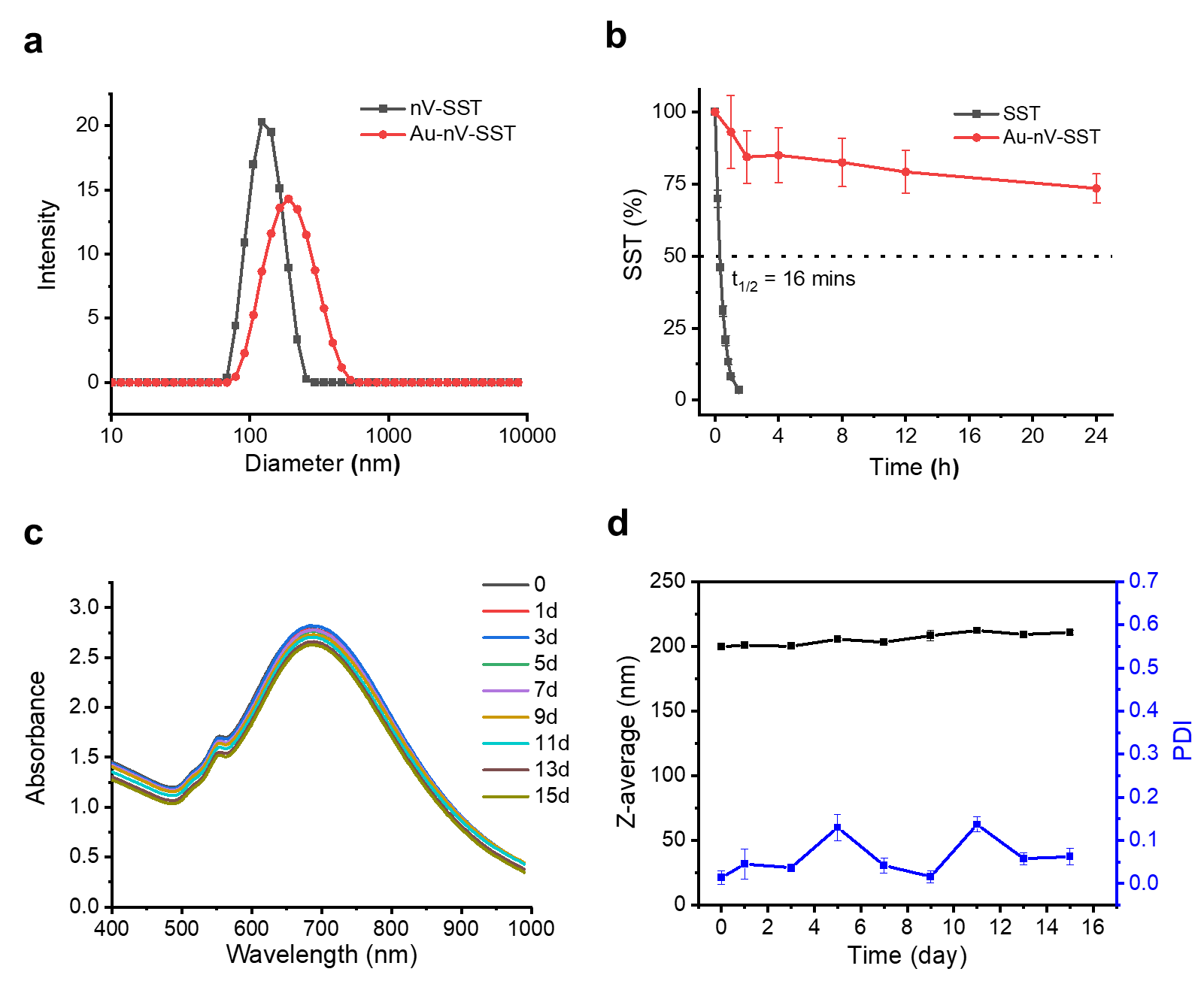

**Supplementary Fig. 1 Characterization of SST-encapsulated gold-coated nanovesicles (Au-nV-SST)**. **a**, Size distribution of SST-encapsulated nanovesicles (nV-SST) and Au-nV-SST measured by dynamic light scattering. **b**, Degradation kinetics of SST by α-chymotrypsin in the sample of free SST solution and Au-nV-SST. **c**,**d**, UV-Vis spectra (**c**) and particle size (**d**) changes of Au-nV-SST in PBS at 4 ^o^C over time (n = 3). Data were expressed as Mean ± S.D.

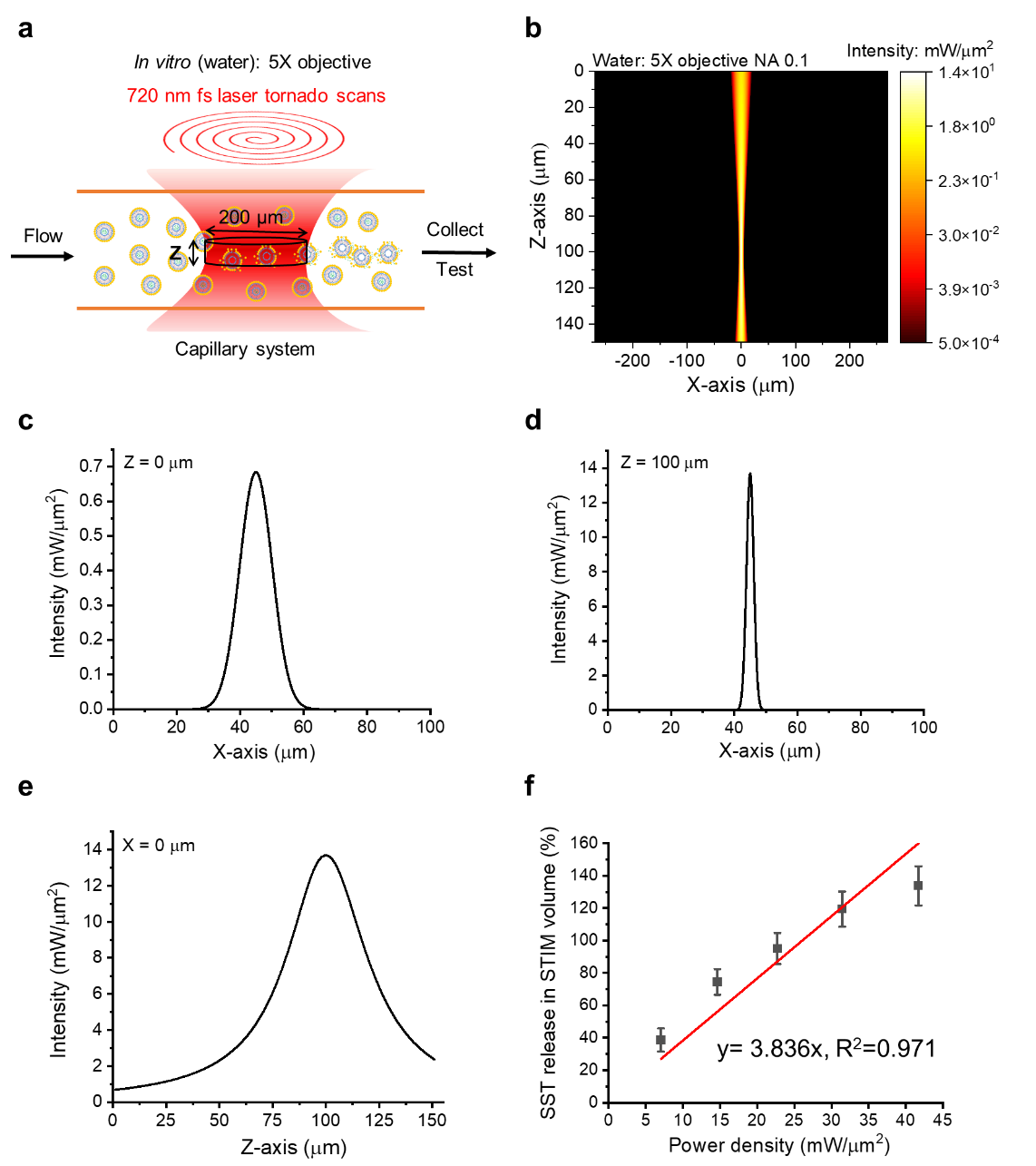

**Supplementary Fig. 2 Numerical analysis on beam propagation of two-photon stimulation *in vitro***. **a**, Schematic of the flow system to measure SST *in vitro* photorelease efficiency (5X objective). Tornado scans (200 µm diameter) are focused at the center of capillary. Effective stimulation in z-axis is considered between the full width of half maximum (FWHM) of light intensity. **b**, 2D map of light intensity distribution in water based on the numerical model (Input: 720 nm, 100 mW). **c**,**d**, Lateral (x-y plane) light intensity distribution at (**c**) the surface (z=0 µm) and (**d**) focus plane (z=100 µm). **e**, Axial light intensity distribution at x=0 µm. **f**, The linear fitting of SST release efficiency in the effective stimulation volume at different power density (720 nm, 40 scans, n =3, raw data from Fig. 1**d**).

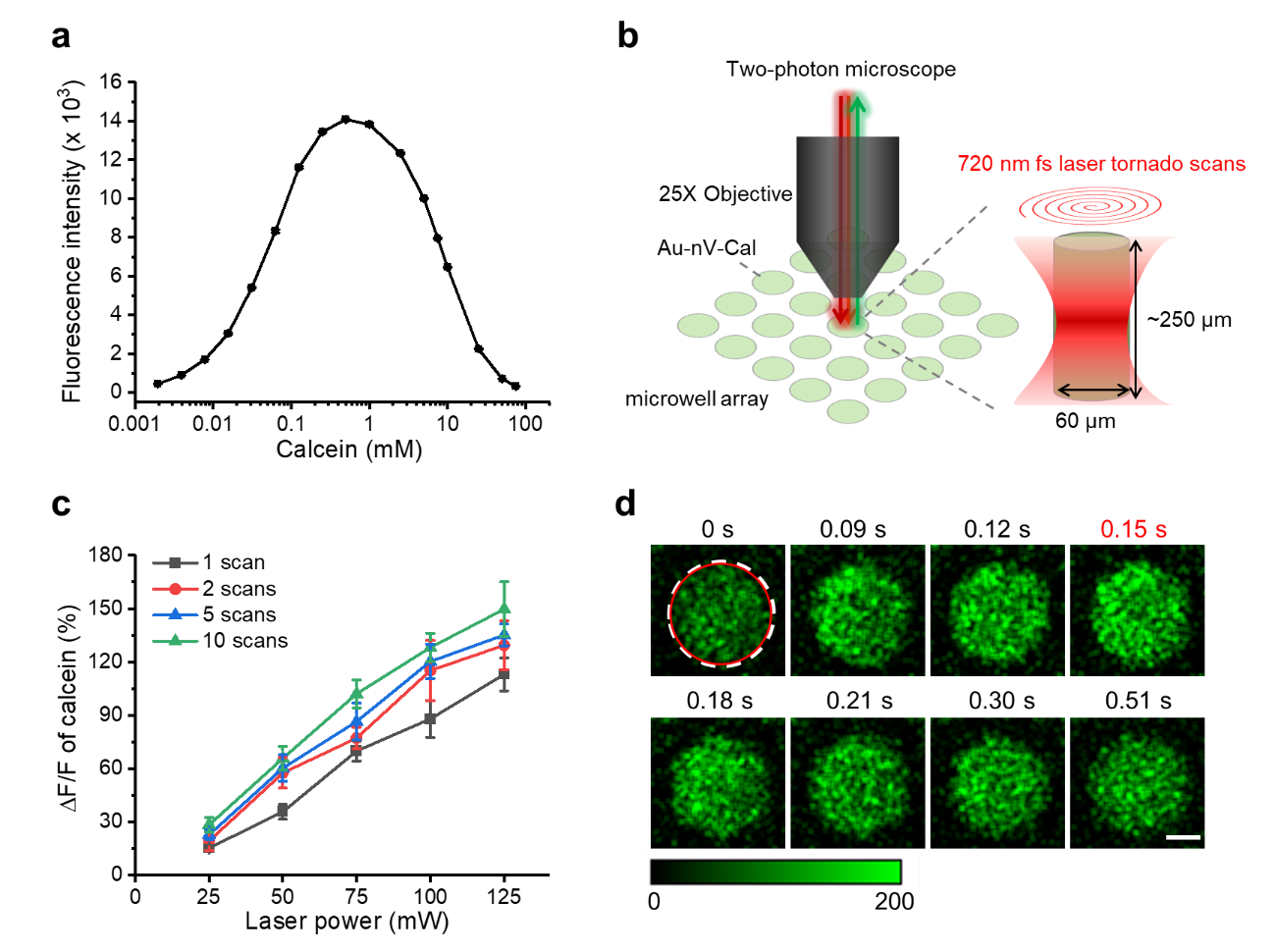

**Supplementary Fig. 3 Calcein release from gold-coated nanovesicles (Au-nV-Cal) under stimulation laser pulses at 720 nm**. **a**, Calcein fluorescence intensity as a function of concentration. Due to the self-quenching, the fluorescence intensity decreases when the concentration is over 0.5 mM. 75 mM of calcein was used to encapsulated in the liposomes. **b**, Schematic of two-photo imaging and stimulation of Au-nV-Cal in the microwell array. Emission of calcein from 495-540 nm was collected at the excitation of 920 nm. Tornado scans (720 nm, 60 µm diameter) are focused at the center of single microwell (60 µm diameter). **c**, Calcein release efficiency at different stimulation power and tornado scans (n = 5). **d**, Real-time fluorescent images of Au-nV-Cal in a microwell (diameter: 60 μm, white dash line). Au-nV-Cal were stimulated at 0 s by a tornado scan (diameter: 60 μm, red solid line) for 65 ms. Scale bar: 20 μm. Data were expressed as Mean ± S.D.

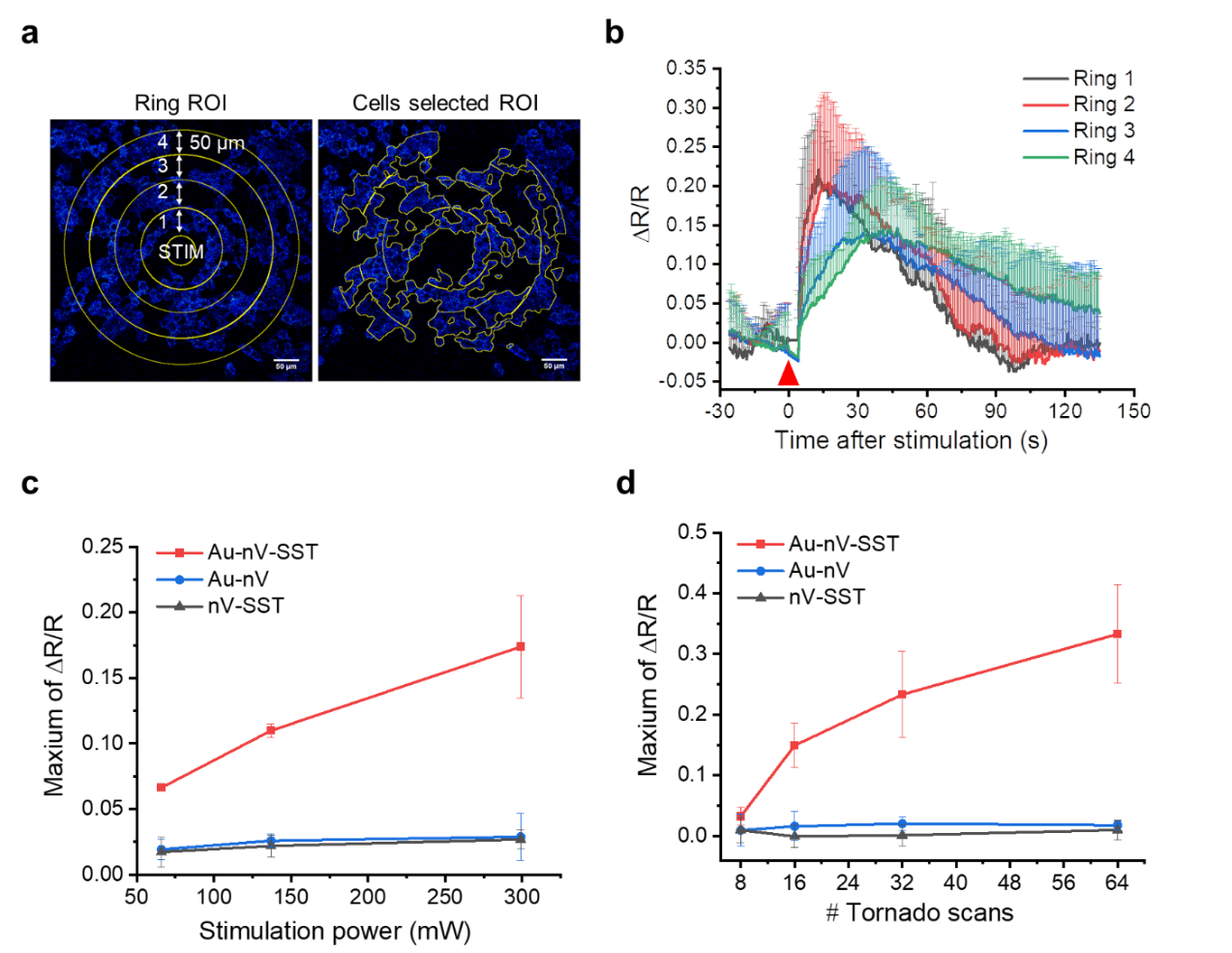

**Supplementary Fig. 4 *In vitro* SST2 CNiFERs response on photoreleased SST**. **a**, Schematic of ring analysis. Tornado scans with a diameter of 60 µm was performed in the center at 0 s. Rings with 50 µm width were created around the stimulation area. Cell were selected in the rings for FRET analysis. Scale bar: 50 µm. **b**, CNiFERs response in different rings (n=4). Ring1-4 correspond to the label in (**a**). Photostimulation was performed at 300 mW for 4 s (64 tornado scans). **c**,**d**, Maximum of CNiFERs response (ΔR/R) as a function of (**c**) stimulation power and (**d**) scans number (n=3). Tornado scans with a diameter of 60 µm was performed besides CNiFERs, and cells in a circle of 200 µm were selected for analysis. Data expressed as Mean ± S.D.

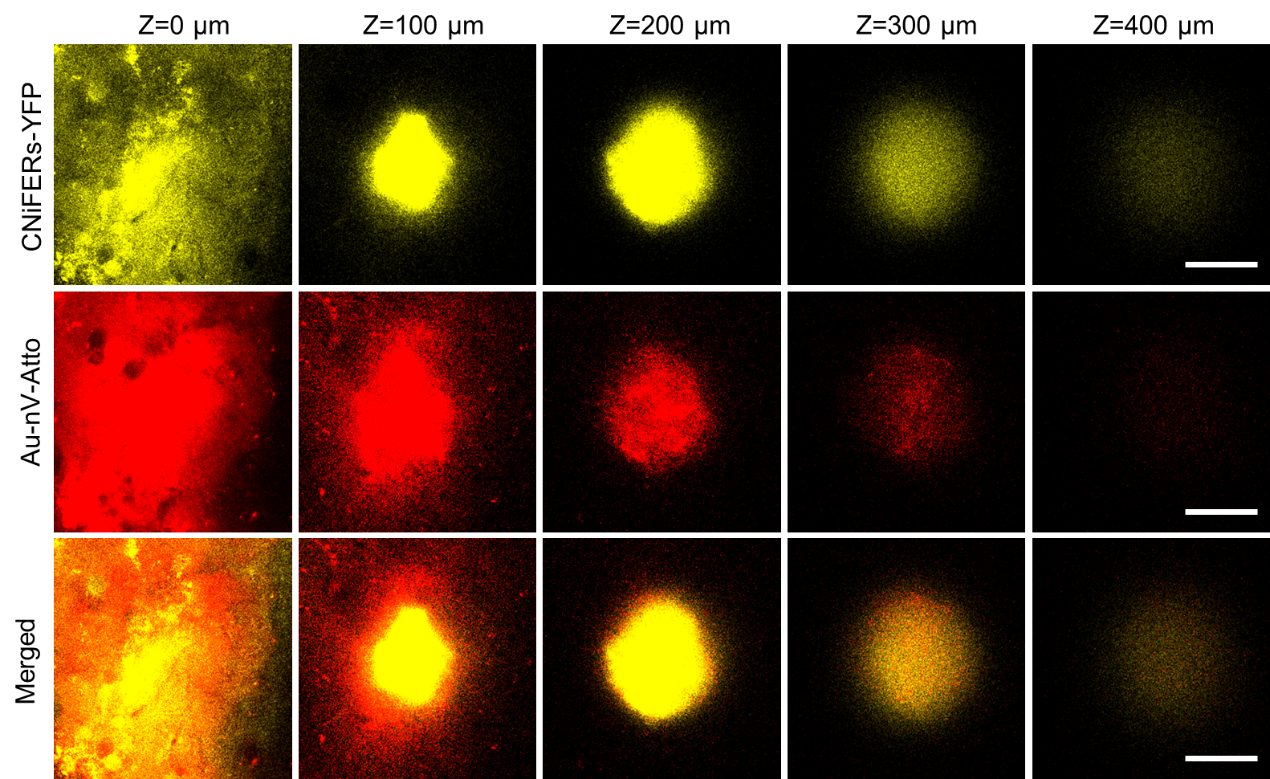

**Supplementary Fig. 5 Z-stacked two-photon fluorescent images of SST2 CNiFERs/Au-nV-Atto in mouse cortex.** Emission of SST2 CNiFERs from 520-560 nm was recorded at the excitation of 900 nm. Emission of Au-nV-Atto from 575-645 nm was recorded at the excitation of 1100 nm. Images were taken from the surface of brain to 400 µm depth. Scale bar: 50 µm.

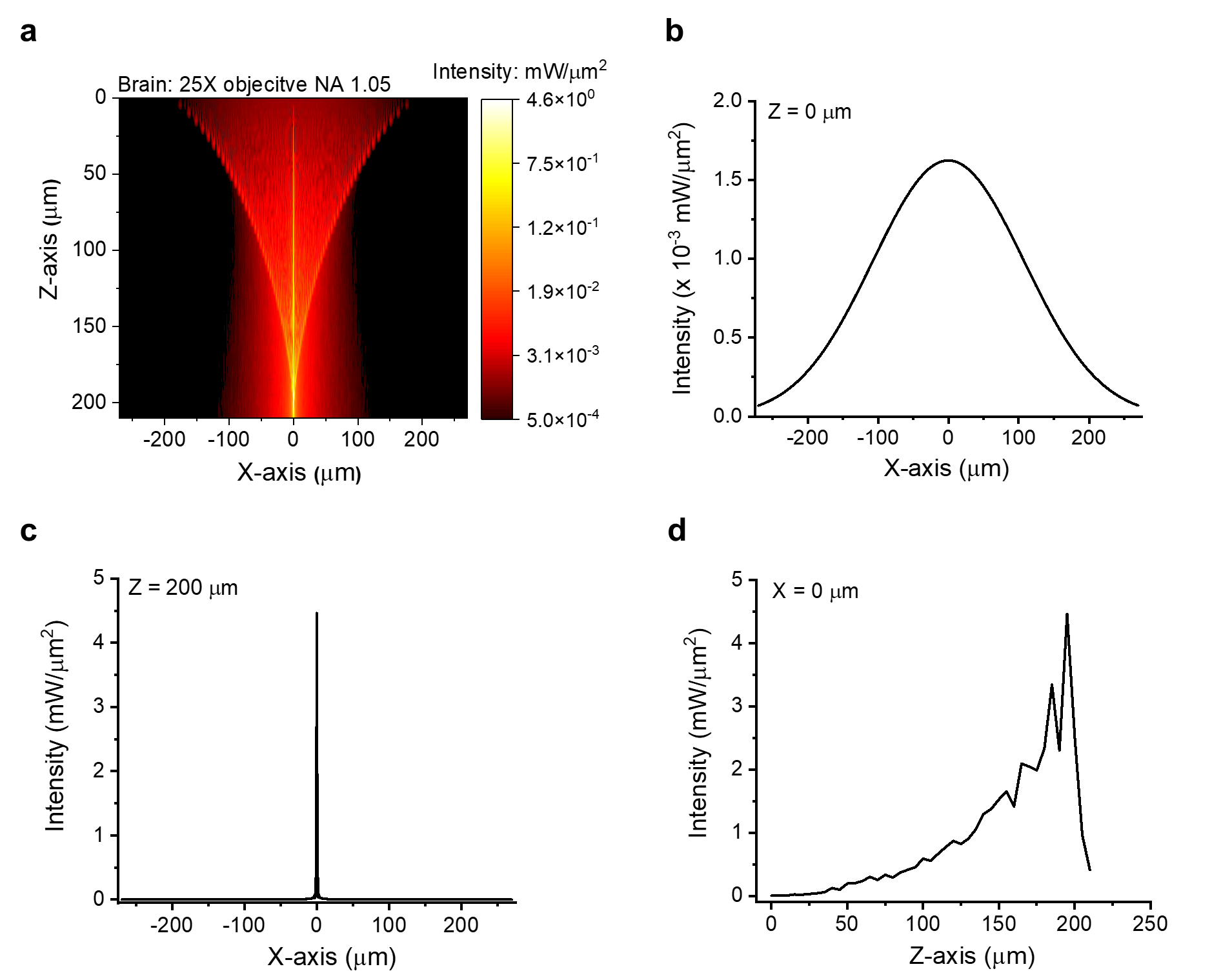

**Supplementary Fig. 6 Numerical analysis of beam propagation of two-photon stimulation *in vivo***. **a**, 2D map of light intensity distribution in mouse brain based on the numerical model (Input: 720 nm, 100 mW). Photostimulation is focus at 200 µm depth, and effective stimulation in z-axis is considered between the FWHM of light intensity. **b**,**c**, Lateral (x-y plane) light intensity distribution at (**b**) the surface (z=0 µm) and (**c**) focus plane (z=200 µm). **d**, Axial light intensity distribution at x=0 µm.

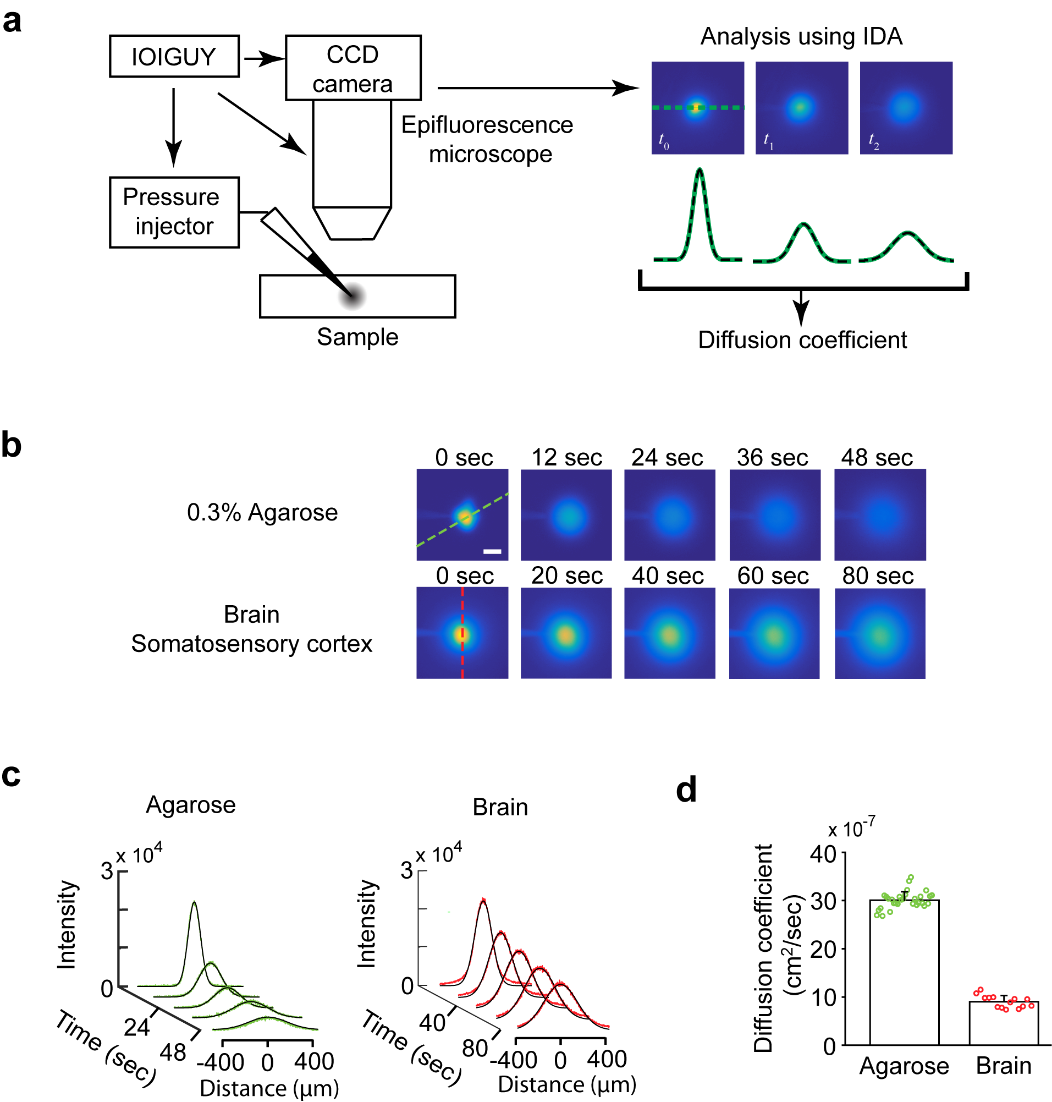

**Supplementary Fig. 7 Quantification of diffusion coefficient of fluorescein-5(6)-carbonyl- somatostatin-14 (5(6)-FAM-SST) using integrative optical imaging (IOI).** **a**, Schematic of the setup and procedure of IOI technique. A small quantity of 5(6)-FAM-SST in PBS was injected into the sample (0.3% dilute agarose gel or acute brain slice) from a microelectrode using a brief pressure pulse. Then, series of images were taken to capture the diffusion by a charged-couple device (CCD) camera attached onto an epifluorescence microscope. The steps of pressure injection and imaging were controlled automatically using a customized MATLAB-based program (IOIGUY). Diffusion coefficient was obtained after the intensity profiles along an axis on the raw images were fitted to the diffusion equation using another customized MATLAB-based (IDA). **b**, Representative fluorescent images of the diffusion of 5(6)-FAM-SST in agarose gel and in brain, respectively. Scale bar: 100 μm. **c**, Series of intensity profiles obtained from raw images. The raw data is shown in color and the fits are shown in black. **d**, Average diffusion coefficients of 5(6)-FAM-SST in dilute agarose gel and in brain. The error bars show the standard deviation, while the circles show individual records in both groups. All values are at 31°C. Free diffusion coefficient (D) = (3.00 ± 0.18) × 10^-6^ cm^2^•s^-1^ (n=30 records); effective diffusion coefficient (D*) = (8.9 ± 1.3) × 10^-7^ cm^2^•s^-1^ (n=14 slices from 5 mice). Data were expressed as Mean ± S.D.

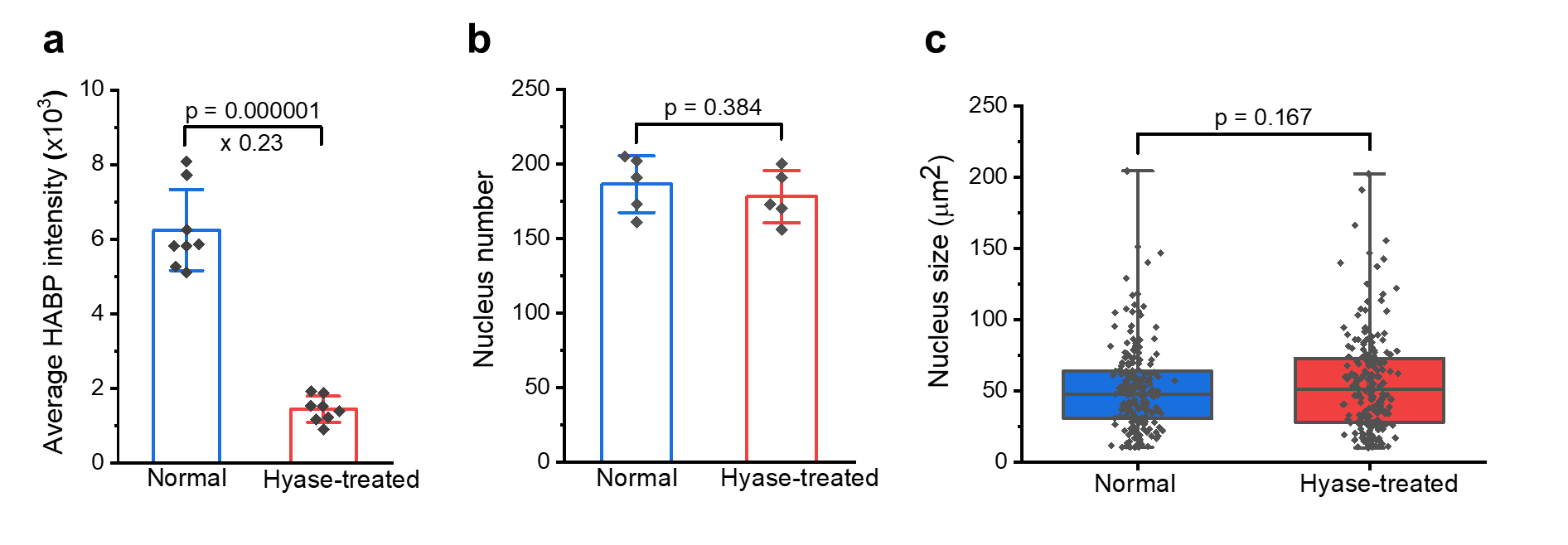

**Supplementary Fig. 8 Quantitative analysis of HABP-labelled normal mouse brain and hyase-treated brain sections. a**, Average fluorescence intensity for HABP in mouse cortex (0.8 mm x 0.8 mm) in both conditions (n = 8 slices). **b**,**c**, Comparison of (**b**) nucleus number and (**c**) nucleus size in the cortex (0.8 mm × 0.8 mm) of normal mouse and hyase-treated mouse brains (n = 5 slices). Data were expressed as Mean ± S.D. in (**a**). For the Box plot (**b**), the lower outlier: minimum; lower line: first quartile; center line, median; upper line: third quartile; upper outlier: maximum; center blank square: mean. Two-sample Student’s t-test in Origin 2021 software was conducted to determine p values and statistical significance.

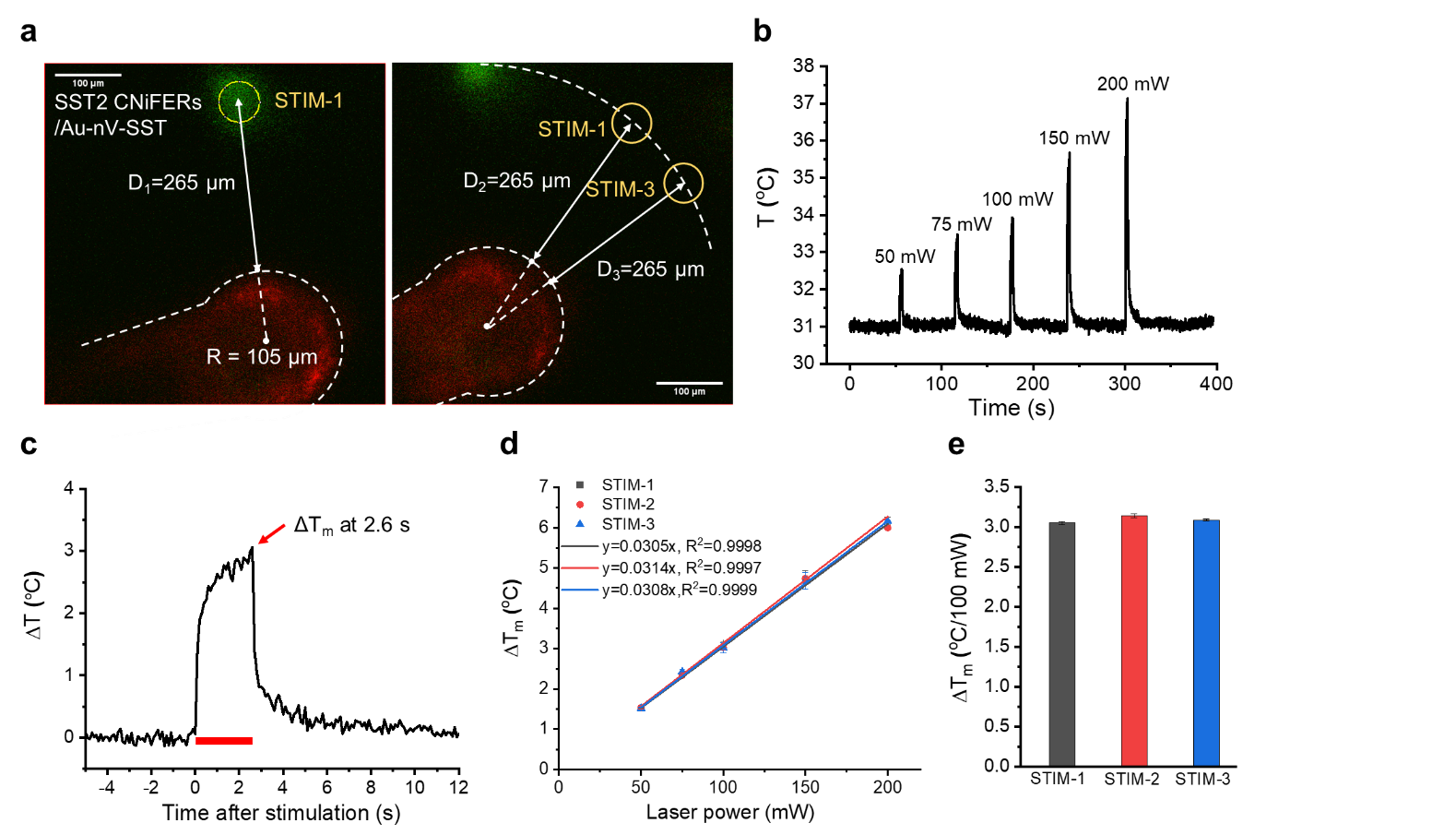

**Supplementary Fig. 9 Temperature change in the mouse brain under two-photon stimulation. a**, Two-photon fluorescent images of SST2 CNiFERs/Au-nV-SST implant (Green, E_m_: 495-540 nm) and quantum dot-coated thermocouple probe (Red, E_m_: 575-645 nm) when excited at 900 nm at 200 µm depth in mouse cortex. The distance from the center of stimulation to the edge of probe is 265 µm. Scale bar: 100 µm. **b**, Real-time temperature recording of mouse brain at 200 µm depth (10 counts/s). The peaks show that the temperature increased when stimulating on the implant (STIM-1) at different power (40 scans, duration: 2.6 s). **c**, The zoomed-in kinetics of temperature increase (ΔT) upon the stimulation of 100 mW and 40 scans (2.6 s). ΔT quickly increased to 1.4 ^o^C in 0.1 s and slowly increased to the maximal value 3 ^o^C at 2.6 s during stimulation. After stimulation, ΔT quickly decreased to 1.4 ^o^C in 0.1 s and slowly decreased to 0 in several seconds. **d**, The linear fitting of maximal temperature increase (ΔT_m_) at different locations and laser power (n=3). STIM-1, STIM-2 and STIM-3 were from the same distance to the probe. No implants were in STIM-1 and STIM-2 areas. **e**, ΔT_m_ per 100 mW acquired from the fitting in (**d**) at different locations. Data were expressed as Mean ± S.D.

**Supplementary Table 1. Laser beam propagation of two-photon stimulation**

| Parameters | 5x objective (*in vitro*) | 25x objective (*in vivo*) |
| --- | --- | --- |
| Input power (mW) | 100 | 100 |
| Numerical aperture (NA) | 0.1 | 1.05 |
| Scattering mean free path (μm) | 20000000 | 122 |
| Wavelength (nm) | 720 | 720 |
| Focus plane depth (μm) | 100 | 200 |
| Fluorescence-E relation | ∝‖E(x,y,z)‖^2^ | ∝‖E(x,y,z)‖^2^ |
| FWHM of Z-axial (μm) | 45.7 | 11.3 |
| FWHM of X-axial (μm) | 2.7 | 0.93 |
| Effective power density (mW/µm^2^)^a^ | 9.5 | 3.6 |
| Laser stimulation volume V_stim_ (nL)^b^ | 2 | 0.032 |
| Release efficiency for one stimulation (40 scans) (α, %)^c^ | 35.6 | 13.8 |

Note: a, average power density above the FWHM (full width of half maximum);

b, Effective stimulation volume when average power density is above the FWHM. For *in vivo*, V_stim_=π·(60µm)^2^/4·11.3µm=0.032 nL

c, acquired from Figure S4f when x=9.5 and 3.6 mW/µm^2^ respectively.

**Supplementary Table 2. Estimation of total released SST number upon photostimulation *in vivo***

|  | Parameters | Methods or formula | Number |
| --- | --- | --- | --- |
| Au-nV-SST | Encapsulated SST concentration in Au-nV-SST C_sst_ (µM) | ELISA | 1.8 |
| Mixture of SST2 CNiFERs and Au-nV-SST | Volume of Au-nV-SST solution per 10^6^ cells V_1_ (μL) | - | 1 |
|  | Diameter of HEK293 cell D_cell_ (μm) | Ref^1^ | 14 |
|  | Volume of 10^6^ cells V_2_ (μL) | π/6·(D_cell_)^3^·10^6^ | 1.4 |
|  | SST concentration in the mixture C_sst_′ (µM) | C_sst_·V_1_/(V_1_+V_2_) | 0.74 |
| SST2 CNiFERs and Au-nV-SST implant | Inject volume V_i_ (nL) | - | 120 |
|  | Average diameter of CNiFERs implant D_c_ (μm) | Measurement | 80 |
|  | Average height of CNiFERs implant h_c_ (μm) | Measurement | 400 |
|  | Volume of CNiFERs implant in brain V_c_ (nL) | π/4·(D_c_)^2^·h_c_ | 2.0 |
|  | Volume of nanovesicles implanted V_n_ (nL) | V_i_-V_c_ | 118 |
|  | Total number of SST injected Q_t_ (X10^10^) | C_sst_’·V_n_·N_A_ | 5.3 |
| Stimulation | Diameter of stimulation D_s_ (μm) | - | 60 |
|  | FWHM of Z-axial h_s_ (μm) | Table 1 | 11.3 |
|  | Laser stimulation volume in brain V_stim_ (pL) | π/4·(D_s_)^2^·h_s_ | 32 |
|  | Stimulated volume ratio R_s_ (%) | V_stim_/V_total_ | 1.6 |
| Photoreleased SST | Release efficiency for one stimulation α (%) | Table 1 | 13.8 |
|  | Number of released SST for one stimulation Q (X10^8^) | Q_t_·R_s_·α | 1.2 |
|  | Local concentration of photoreleased SST in the CNiFER implant (nM) | Q//(N_A_⋅V_c_) | 100 |
